# Fast leaps between millisecond confinements govern Ase1 diffusion along microtubules

**DOI:** 10.1101/2021.04.15.439939

**Authors:** Łukasz Bujak, Kristýna Holanová, Antonio García Marín, Verena Henrichs, Ivan Barvík, Marcus Braun, Zdenĕk Lánský, Marek Piliarik

## Abstract

Diffusion is the most fundamental mode of protein translocation within cells. Confined diffusion of proteins along the electrostatic potential constituted by the surface of microtubules, although modeled meticulously in molecular dynamics simulations, has not been experimentally observed in real-time. Here, we used interferometric scattering microscopy to directly visualize the movement of the microtubule-associated protein Ase1 along the microtubule surface at nanometer and microsecond resolution. We resolved millisecond confinements of Ase1 and fast leaps between these positions of dwelling preferentially occurring along the microtubule protofilaments, revealing Ase1’s mode of diffusive translocation along the microtubule’s periodic surface. The derived interaction potential closely matches the tubulin-dimer periodicity and the distribution of the electrostatic potential on the microtubule lattice. We anticipate that mapping the interaction landscapes for different proteins on microtubules, finding plausible energetic barriers of different positioning and heights, will provide valuable insights into regulating the dynamics of essential cytoskeletal processes, such as intracellular cargo trafficking, cell division, and morphogenesis, all of which rely on diffusive translocation of proteins along microtubules.

## Introduction

Microtubules (MTs), one of the key components of the cytoskeleton, are filaments formed by alpha- and beta-tubulin heterodimers assembled head to tail with 8 nm periodicity, forming protofilaments that form a closed hollow tube with ~25 nm diameter. A plethora of proteins interact with MTs to fulfill a variety of cellular processes. In particular, MTs form tracks for translocation of a multitude of these MT-associated proteins (MAPs). MAP translocation along the MT lattice underpins, e.g., long-range transport of intracellular cargo or the integrity of the MT network during vital processes, such as cell division. Enabling the various tasks fulfilled by MAPs, the movement of these proteins along the MT lattice occurs in two main ways. Although some MAPs, including the majority of MT-associated molecular motors, move along the MTs unidirectionally, many MAPs diffuse along the MT lattice performing a random motion driven by thermal energy. Lattice diffusion has been shown to be an efficient mechanism for MAPs to target the MT tips and enable the regulation of the MT assembly and disassembly,^1^ enable crosslinking of MTs into pliable bundles,^2^ or tether molecular motors to the lattice.^3^ Although they have been extensively studied both experimentally and theoretically,^4–9^ the mechanistic understanding of the diffusible motion of MAPs along the MT lattice is still lacking, mainly owing to the insufficient spatiotemporal resolution of the employed experimental techniques.

Microtubule crosslinker Ase1 (anaphase spindle elongation 1), a member of the Ase1/PRC1/MAP65 family, is a prototypical diffusible MAP. Ase1 is essential during cell division^10–12^ owing to its ability to bundle MTs and stabilize MT networks^13^ through the generation of entropic and frictional forces.^2,5,14^ The essential feature for this stabilization is a rather low unbinding rate of Ase1 from the microtubule, combined with its ability to rapidly diffuse along the microtubule surface (diffusion coefficient ~ 0.1μm^2^ s^-1^) over timescales of several minutes^5,15^. The interaction of Ase1 with MTs is mediated by the Ase1 spectrin domain and an unstructured C-terminus.^6,7^ The mechanism that allows the Ase1 molecule to stay in contact with the MT for prolonged periods but still allows for rapid diffusion along the MTs is still not fully understood. Theoretical studies indicate the central role of electrostatic interactions in protein diffusion characterized by discreet steps of dimeric tubulin.^9^ Despite numerous hypotheses,^9,16^ it remains unclear whether the diffusibility is owing to transient unbinding and rebinding, leading to “hopping” along with the periodic microtubule-binding potential, or whether Ase1 can “slide” within a relatively shallow microtubule potential.

The motion of single Ase1 and other diffusible molecules along the microtubule has mostly been studied using total internal reflection fluorescence (TIRF) microscopy and optical tweezers. Optical tweezers can be used to probe the single-protein interaction by applying an external load, whereas direct observation of the single-protein trajectory through microscopy can be used to explore its interaction driven only by the thermal fluctuations of the environment. However, single-molecule fluorescence microscopy has several technical and fundamental limits. Processes such as saturation, photobleaching, and blinking^17,18^ limit the number of photons collected from a single fluorophore at a given time scale and consequently hamper the temporal resolution as well as localization precision of the single emitter. To overcome those limitations, scattering labels such as small metallic or dielectric nanoparticles or even small biomolecules can be directly detected by using elastic scattering^19^. Dark-field microscopy has been used to demonstrate single nanometer precision of localization to visualize the rotation of F_1_-ATPase^20^ and to describe previously unresolved fine details of the kinesin stepping motion.^21^ Interferometric scattering microscopy (iSCAT) has enabled the tracking of nanoparticles as small as 2 nm at high spatial and temporal resolution.^22,23^ It has been used to observe the motion of motor proteins,^24,25^ the interactions of membrane proteins,^26^ protein diffusion in lipid membranes and live cells,^27,28^ label-free detection of single proteins,^29,30^ and protein conformational changes.^31^

Here, we explore fine details of the interaction of the Ase1 protein with the MT lattice by employing fast 3D tracking of single proteins with nanometer precision and 20-μs temporal resolution. Within the Ase1 trajectories, we reveal millisecond and sub-millisecond confinements associated with the Ase1 interaction with single-tubulin dimers. We were able to resolve the stepping character of the diffusion and extract the spatiotemporal stepping statistics of the Ase1 motion. The achieved measurement accuracy and the strict periodicity of the tubulin lattice allowed us to extract the profile of the periodic interaction potential of Ase1 with the surface of the microtubule. We visualized the distribution of the interaction potential of a single tubulin dimer and complemented the experimental data with corresponding molecular dynamics (MD) simulations, which closely matched the quantitative values derived from the experiment.

## Results

### High-fidelity tracking of Ase1 diffusion on a microtubule

To study the interaction between Ase1 and an MT, we used a custom-built interferometric scattering microscopy (iSCAT) setup described in the Methods. This instrumentation enabled the imaging of the position of unlabeled MTs and tracking the position of Ase1 labeled with a scattering label with nanometer precision in all three coordinates. We immobilized double-stabilized MTs through electrostatic interaction to the APTES-coated microscope coverslips and added 0.60 nM His-tagged Ase1 construct in the motility buffer (Methods). To visualize the Ase1 trajectory (Figure 1b), we specifically attached gold nanoparticles (GNP) to single Ase1 molecules inside the flow chamber (Supplementary Fig. S8). The Ni-NTA functionalization protocol ensured that the vast majority of GNP labels were attached to exactly one Ase1 molecule (Methods, Supplementary Fig.S4).

**Figure 1.**
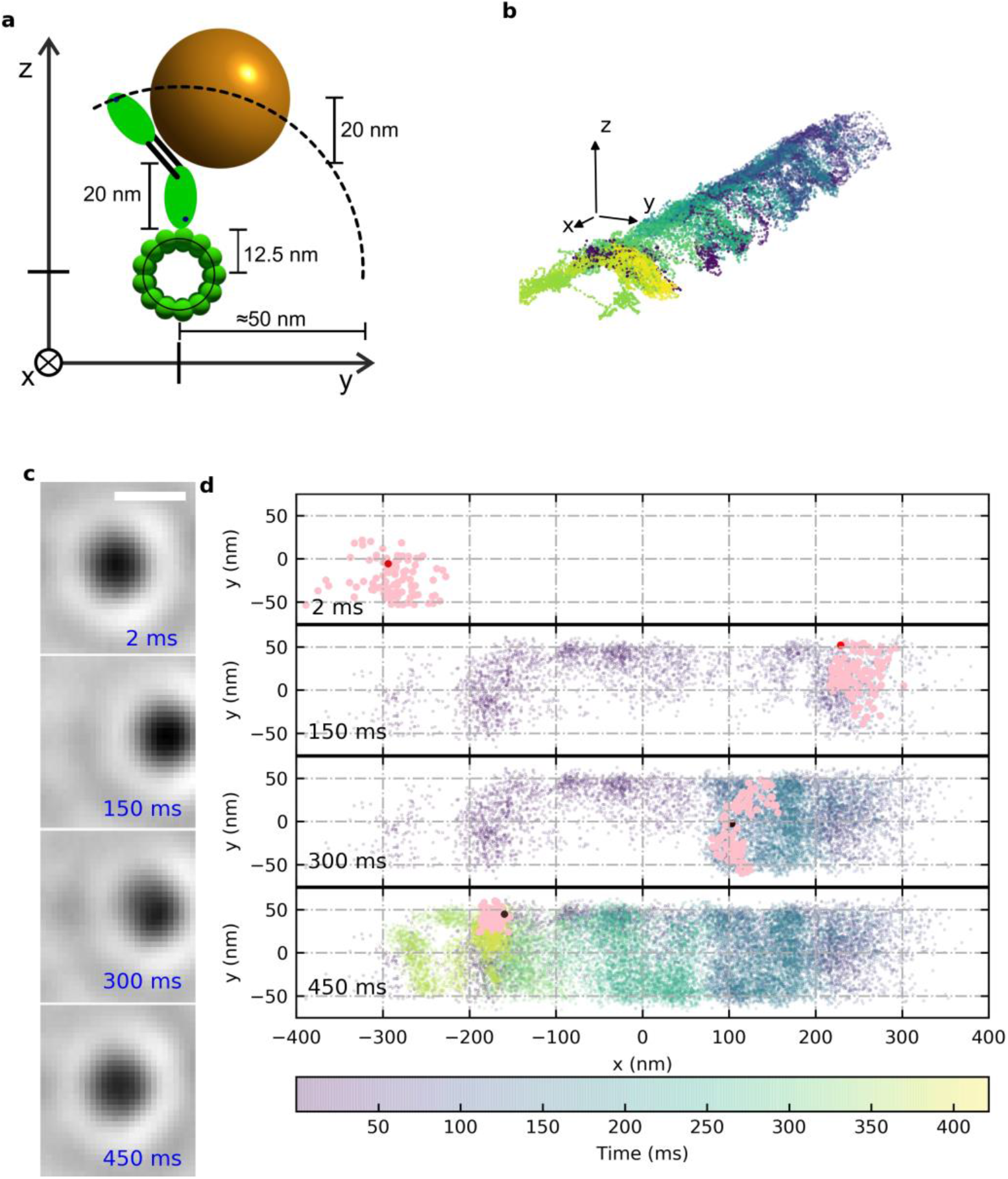
Measurement and localization of single Ase1 protein. **a,** Scheme of the experimental system consisting of a microtubule, Ase1 protein, and 40-nm GNP attached to the dimerization domain of the Ase1. **b,** A 3D visualization of the Ase1 motion on MT. Scale bars in all directions = 20 nm. **c,** Sequence of background-corrected iSCAT images of labeled Ase1 at selected time stamps of the experiment. Scale bar = 500 nm. **d,** The time-resolved visualization of the 45 kHz trajectory of the Ase1-GNP construct. The last 2 ms of the trajectory at selected time-stamps are indicated in pink. Time is color-coded in (b) and (d).

The position of the GNP-labeled Ase1 was imaged and tracked at 45 000 frames per second (fps) and an example set of iSCAT images after referencing the static background is included in Figure 1c, each showing a diffraction-limited dip of ~8 pixels full-width-at-half-maxima (FWHM), which corresponds to ~300 nm. During the 500 ms measurement, the protein was diffusively displaced on the MT within a range of ~1200 nm, consistent with the previously reported diffusive behavior of Ase1 on an MT.^4,5,15^ We localized the position of the GNP label in each frame and the corresponding high-resolution trajectory of the protein is shown in Figure 1d, which indicated that the Ase1 molecule diffused not only along one microtubule protofilament but also frequently switched between protofilaments. Therefore, the motion of the Ase1 molecules is better described by a 2D diffusion on the surface of the microtubule. Along the axis perpendicular to the MT, the GNP diffusion approximately renders a ribbon of 100 nm width, which closely matches the range of possible lateral positions of the GNP relative to the MT schematically described in Figure 1a. Combined, our high-resolution data show that Ase1 diffusively explores the whole accessible surface of the MT.

### Extracting the 3D envelope of Ase1 motion

Because of the interferometric character of the iSCAT technique, the contrast of the image directly correlates with the distance between the GNP and the coverslip surface.^32,26^ An example of a typical 2D trajectory of Ase1 on an MT is shown in Figure 2a and the dependence of the contrast changes on the displacement perpendicular to the MT is shown in Figure 2b. The iSCAT contrast of the GNP label varied between 35% and 55% and correlated with its immediate position on the MT. Several methods can be used to quantify the vertical coordinate, including model-based linearization of the contrast-displacement calibration,^33^ templated surface-confined trajectory,^32^ advanced fitting of a model point-spread-function^34^ to obtain a fully quantitative phase image of the nanoscatterer.^35^ Considering the narrow range of vertical displacements and simple cylindrical geometry of the system, we used a combination of the first two approaches. We mapped the contrast variation linearly on the vertical displacement and adjusted the scaling factor to preserve the expected cylindrical symmetry of the trajectory template, as shown in Figure 2b, and visualized the reconstructed 3D trajectory (Figure 2d and Supplementary Video S1). The uncertainty in the radial coordinate *ρ_NP_* is not limited by the localization precision in either direction but rather a spherically symmetric fluctuation in the particle position. The localization precision of the GNP label was between 0.5 nm and 1 nm, both in the lateral x-y coordinates and in the vertical z coordinate (Supplementary Fig. S1), whereas the fluctuation of the Ase1-linked GNP rendered a spherically symmetrical volume of the standard deviation of 2.1 nm diameter (Supplementary Information and Supplementary Fig. S2).

**Figure 2.**
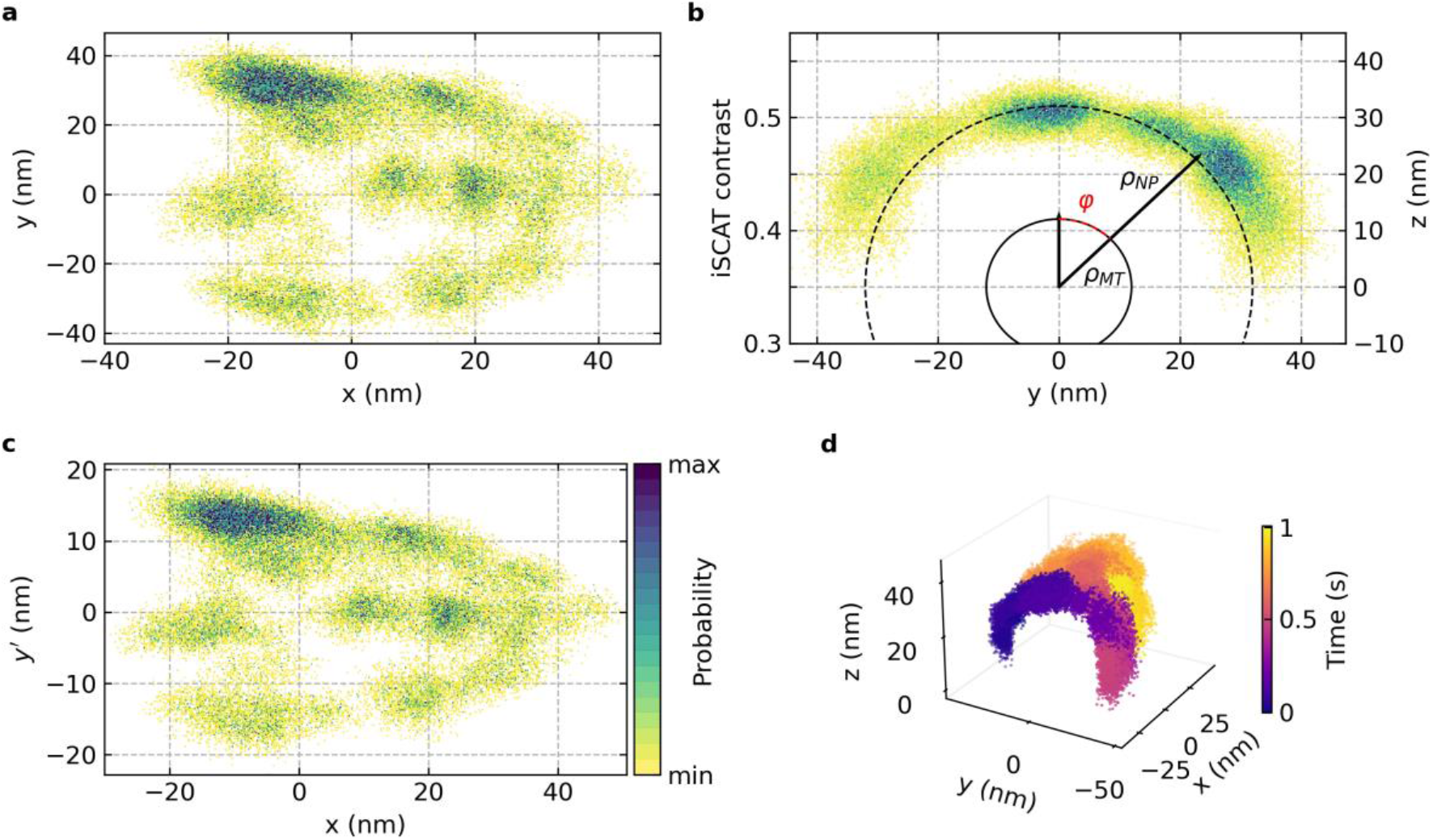
Unwrapping the cylindrical diffusive trajectory of a single protein. **a,** Selected trajectory of the Ase1-GNP construct as detected from the localization of the GNP position. **b,** Dependence of the contrast on the displacement perpendicular to the MT axis. Z position (right axis) is deduced from contrast fluctuations. The dashed line indicates the position of the GNP label and the solid circle indicates the expected position of the MT surface. Cylindrical coordinates are indicated in the plot. **c,** Unwrapped trajectory projected on the MT surface. The density of the localization probability is color-coded in (a-c). **d,** A 3D visualization of the trajectory with a color-coded time axis.

Finally, we used the cylindrical coordinate system to unwrap the 3D trajectory into two dimensions, corresponding to the longitudinal coordinate *x* and the transverse displacement *y*′ = *ρ_MT_φ* of the nanoparticle projected to the MT surface (Figure 2c).

### Decoupling transversal and longitudinal diffusion

We collected an extended dataset (n = 83) of different Ase1 trajectories on the surface of the MTs under identical experimental conditions. Four rather diverse examples of the resulting trajectories spanning from the slowest/least diffusive examples to the fastest/most diffusive ones are shown in Figure 3a-d. Trajectories were corrected for the cylindrical motion and unwrapped along the radius of the MT. All trajectories clearly show a two-dimensional (2D) diffusion pattern confined within the diameter of the microtubule.

**Figure 3.**
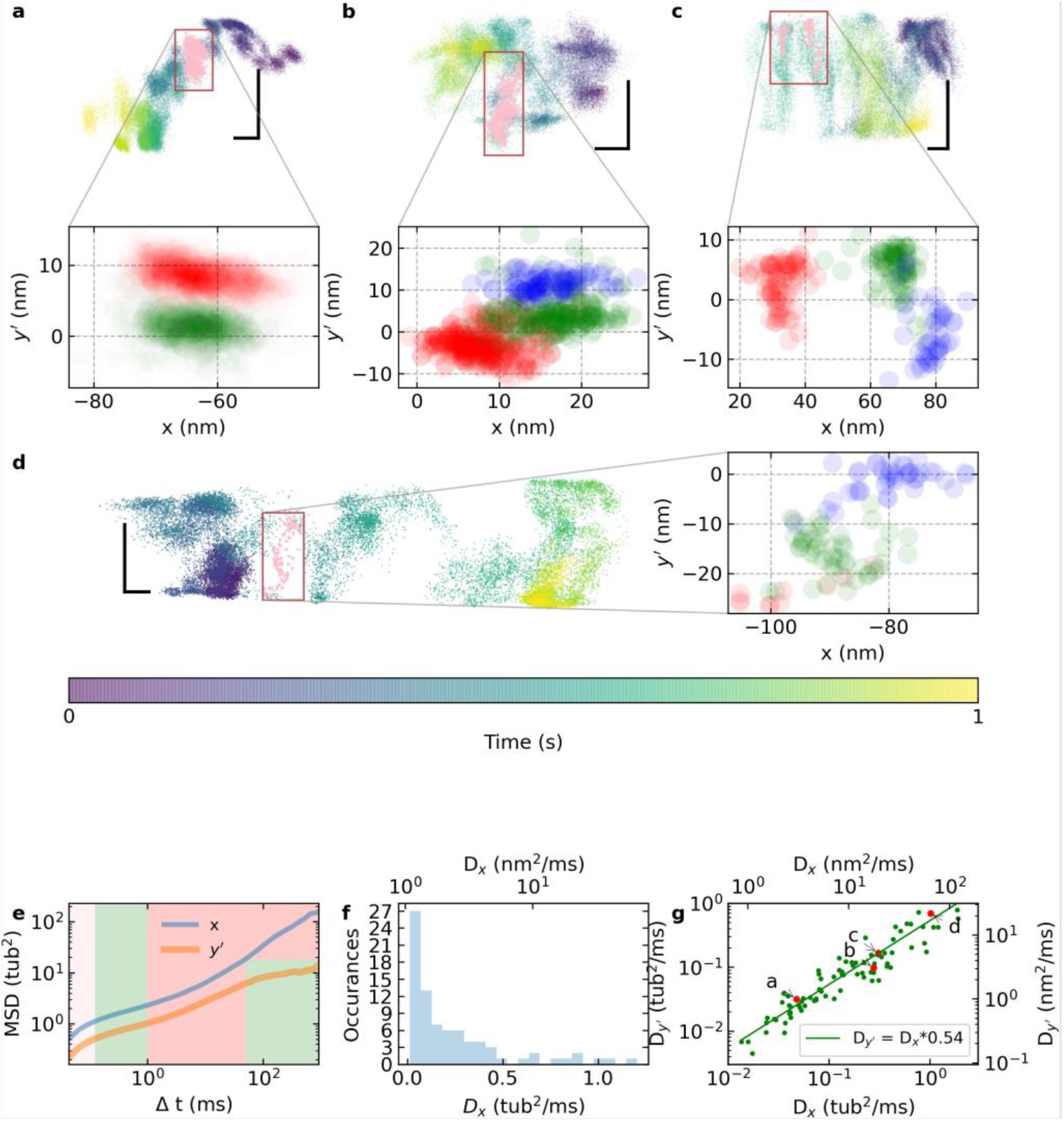
Diffusive behavior of Ase1 on the MT surface. **a-d,** One-second Ase1 trajectories collected at 45 kHz framerate with a color-coded time axis. Horizontal and vertical scale bars are 25 nm. The zoomed-in sections show segments of the trajectory highlighted in pink. Colors discern the time sequence of confinements. Note that in (b) and (c) the particle travels multiple times through the cropped area and only one pass is highlighted. **e,** Average of the longitudinal (x) and transverse (y’) MSD plots (all 83 MSD plots are shown in Supplementary Fig. S5a-b). Green regions indicate confined diffusion and red regions highlight the interval of close-to-free diffusion. **f,** Histogram of longitudinal diffusion coefficients along the x-direction. **g,** Correlation of the transverse and longitudinal diffusivities. Red points show the specific cases selected in a-d.

We analyzed all the measured Ase1 trajectories using the mean square displacement (MSD) statistics separately along the parallel and perpendicular coordinates. The collection of all the MSD plots is provided in Supplementary Fig. S5a-b. The MSD curve averaged from all the individual trajectories in the longitudinal and the transverse axis are plotted in Figure 3e. Both the transversal and the longitudinal MSD plots start with an ascending slope on the sub-100 μs timescale. The absence of a plateau at the shortest time intervals indicates that the single-frame displacement (averaged at 3 nm) is not limited by the localization precision of the experiment but reflects the diffusive motion of the Ase1-GNP construct. The diffusive segment of the MSD plot at the shortest time intervals is followed by a clear plateau between 100 μs and 1 ms, indicating transient confinements^36^ appearing in the vast majority of the trajectories. The underlying details of these confinements are highlighted in the cropped sections in Figure 3a-d (smoothed with a 330 μs window). We hypothesize that these confinements are associated with the interaction of the Ase1 molecule with single tubulin dimers in the MT lattice. For time delays longer than the confinement time, the trajectories revert to their diffusive character and in the case of the transversal component saturate at approximately 50 ms as a result of the spatial confinement to the MT diameter. We first analyzed the long-range diffusive character of the trajectories along the MT axis. All axial MSD plots were fitted with a linear diffusion model (Methods) in the interval of 10 - 100 ms to extract the axial diffusion *D_x_* (Figure 3f). To avoid any ambiguity in comparing the displacement in the transversal and longitudinal direction, we expressed the diffusivity in tubulin^2^ ms^-1^ (1 tub^2^ ms^-1^ ≈ 64 nm^2^ s^-1^ in the longitudinal direction). Despite remaining uncertainty in the particular MT conformation, we calibrated the single tubulin-dimer displacement as 8 nm in the axial direction and 26° in the transversal direction, corresponding to 6 nm on the unwrapped *y’* axis (considering the most probable 14-3 architecture of MT in our experiments^37^). The limited interval of unconstrained transversal diffusion, however, alters the MSD plot corresponding to the diffusive motion and, thus, prevents direct fitting of the transverse diffusivity coefficient. Therefore, we directly calculated the ratio of the transverse and longitudinal diffusivities *D_y′_*/*D_x_* as the mean value of the ratio of the respective MSDs (Supplementary Fig. S5c-d, Methods) and used it to estimate the diffusivity *D_y′_* as a function of *D_x_* (Figure 3g). The slope of the linear fit in Figure 4c corresponds to a mean *D_y′_*/*D_x_* ratio of 0.54, indicating that the Ase1 molecule diffuses approximately two times faster along the axis of the MT than in the perpendicular direction.

**Figure 4.**
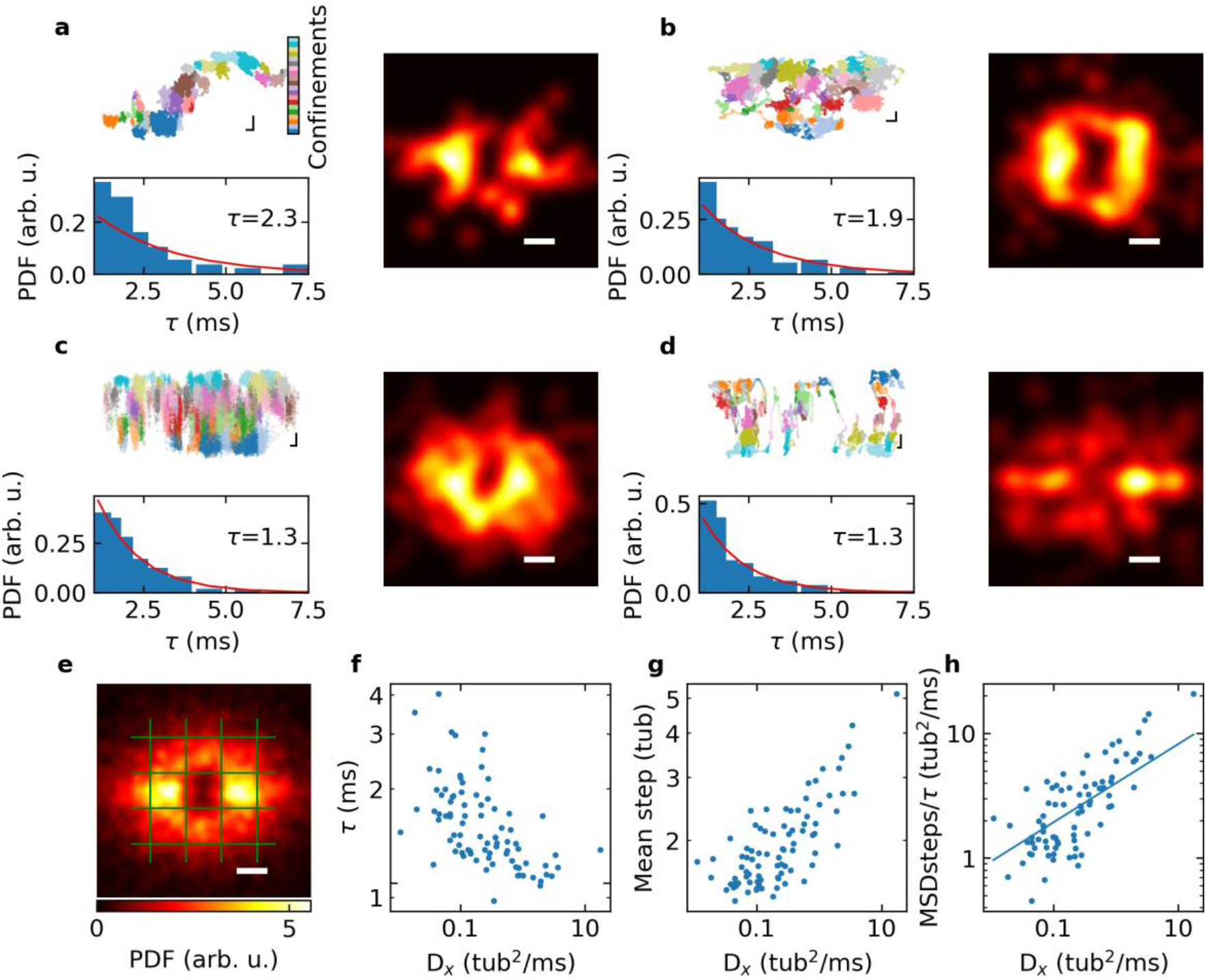
Extracting steps between tubulin dimers. **a, b, c, d** show trajectories of Ase1 with color-coded detected confinements. The step detection analysis shows a histogram of the dwell time in confinements (PDF – power density function of *τ* below each trajectory, fitted mean dwell time in milliseconds indicated in each plot) and a 2D histogram of displacement vector between individual confinements is shown on the right of each trajectory. Scale bars correspond to the size of a single-tubulin dimer. The color scale is indicated by the colorbar in e. **e,** Accumulated 2D histogram of the displacement vector for all measured trajectories. The green grid indicates the position of tubulin dimers in the MT lattice. **f,** The dependence of the mean dwell time of confinements on the longitudinal diffusivity. **g,** The dependence of the mean step length between confinements on the longitudinal diffusivity. **h,** The ratio of MSD per one step and the mean dwell time in each step plotted against the longitudinal diffusivity. The straight line indicates the linear fit crossing the axis origin.

### The stepping character of the diffusion

As indicated by the MSD analysis in Figure 3e, the Ase1 trajectory shows at least two different types of motion on the surface of the MT: (i) millisecond confinements, likely associated with the transient interaction of Ase1 with a single tubulin dimer and (ii) diffusive transitions between these confinements. We detected the position with locally increased density of particle localization occurrences and indexed individual apparent confinements (Methods). Selected trajectories with color-coded apparent confinements are shown in Figure 4a-d and the analysis of all the trajectories is provided in Supplementary Fig. S6. We extracted the dwell time in each of the confinements and histograms of the dwell times suggest an exponential distribution (Figure 4a-d). The mean dwell time of the Ase1 confinements decreased with increasing diffusivity fitted from the MSD plot (Figure 4f); however, the variation in the dwell time does not fully explain the corresponding variation in the diffusivity.

Next, we extracted the displacement vector between individual confinements detected in each trajectory, and the corresponding 2D histograms are provided in each subplot of Figure 4a-d. Distinct maxima within the rectangle aligned approximately with the square of the closest tubulin neighbors are evident in the 2D histograms (see the accumulated displacement histogram in Figure 4e). With increasing diffusivity, the displacement over two or more tubulins becomes more probable (Figure 4g). The accumulated displacement histogram in Figure 4e further shows two peaks of increased step probability in the horizontal direction corresponding to a forward and backward step by a distance of a single tubulin dimer along the MT protofilament. On average, the Ase1 step along the MT axis was more probable by a factor of 2 (*p_x_* = 0.63) compared with the displacement in the transversal direction (*p_y’_* = 0.37) (Supplementary Fig. S7). This behavior was in agreement with the MSD analysis and was consistent with the faster diffusivity in the direction of the MT axis. Interestingly, the ratio of the MSD per one step and the mean dwell time in each step correlated with the diffusivity calculated from the MSD plot (Figure 4h; Pearson correlation coefficient 0.76). This observation indicated that the diffusive motion of the tracked Ase1 was, to a large extent, determined by the step-by-step interaction of Ase1 with the MT. The residual deviation from the strictly linear dependence, as indicated by the linear fit in Figure 4h, may be associated with the limitation of the step-finding algorithm such as possible missed steps resulting in an overestimation of the mean square displacement per single step or a finite time of a single transition between individual confinements.

The vast majority of trajectories (all listed in Supplementary Fig. S6), including the slowest diffusivities (e.g., Figure 4a), clearly show a peak density of single-step displacements within the nearest one-tubulin perimeter. This observation further supports our claim that a single GNP is linked to the MT through a single Ase1 molecule in the majority of experiments. If this was not true, we consider it very unlikely that multiple Ase1 molecules linked to a single GNP would synchronously step from tubulin to tubulin, giving rise to the single-tubulin displacement peak.

### Modeling of the Ase1-MT lattice interaction

From the previous analysis, it remains unclear why Ase1 is more prone to step along the MT protofilament than in the transversal direction, across protofilaments. We hypothesize that it is related to the energetic landscape of the Ase1 interaction with the lattice of tubulin dimers. To better understand the molecule choreography, we performed a coarse-grain simulation of the Ase1 interacting with the tubulin lattice in the starting scenario sketched in Figure 5a. A coarse-grained Cα-based model of an MT lattice was constructed based on the structure PDB ID 5JCO.^38^ It consisted of 3 protofilaments, each consisting of 2 heterodimeric tubulin molecules (i.e., the lattice includes 3×4 monomeric tubulin proteins). The disordered tails with sequences DSVEGEGEEEGEEY for α tubulin (net charge of −8) and DATADEQGEFEEEGEEDEA for β tubulin (net charge of −11) were added as linear chains to the C-terminal of each tubulin monomer. The template structure PDB ID 5KMG^6^ was used in homology modeling of the diffusing Ase1 protein. Again, several C-terminal amino acids were added to form a short unstructured tail. The resulting coarse-grained Cα-based model consisting of 138 beads covered residues Y363–R500 of Ase1 (net charge +9).

**Figure 5.**
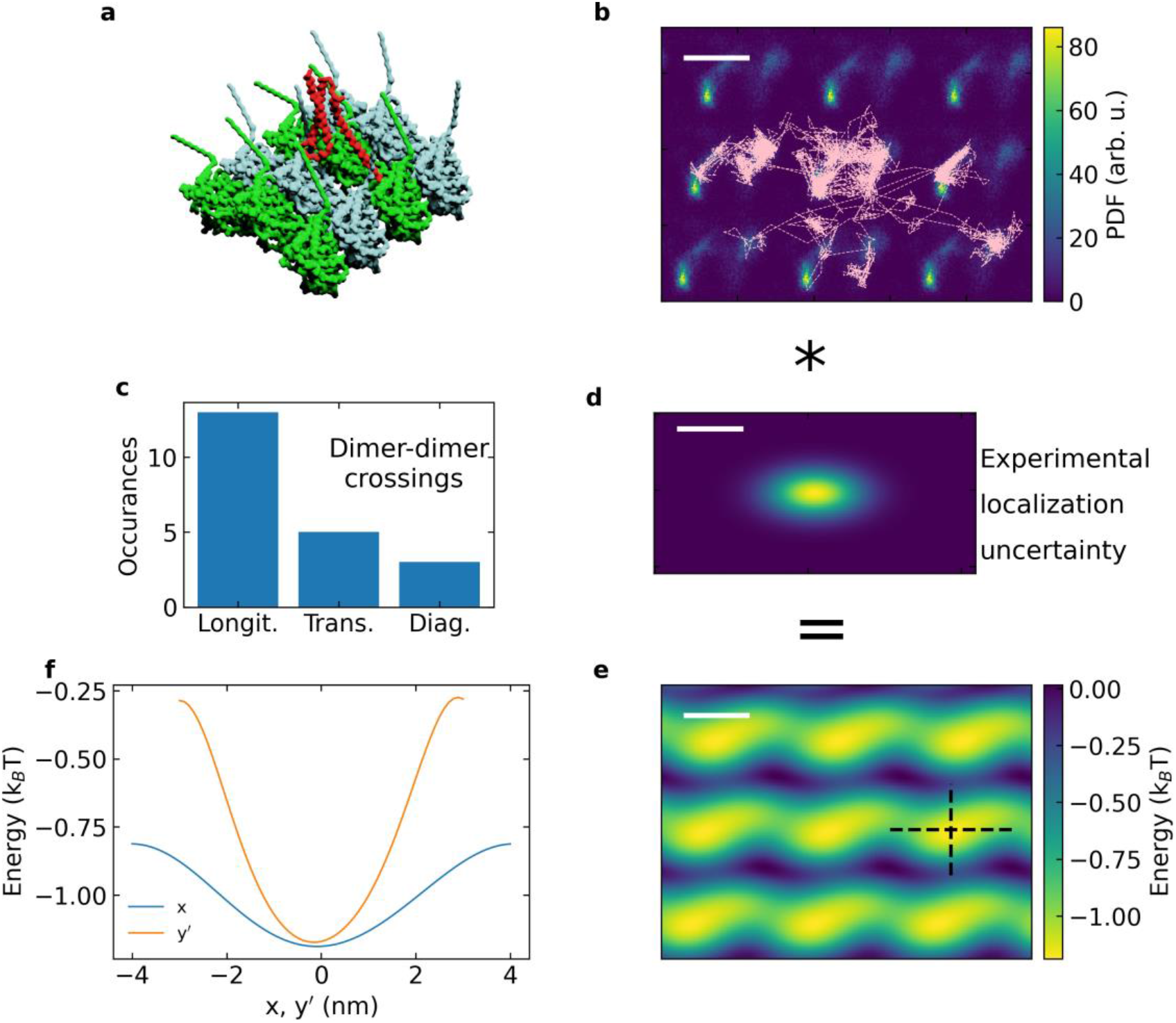
Coarse-grain simulation of the single-tubulin-dimer interaction potential. **a,** Segment of the modeled lattice of tubulin dimers (α- and β-tubulins indicated in green and turquoise, respectively) with the starting position of the segment of the Ase1 molecule (red). **b,** 2D position probability map of the Ase1 molecule with an overlay plot of the simulated set of 13 trajectories (5x binned in time for clarity). **c,** Histogram of observed transitions between tubulin dimers in the longitudinal, transversal, and diagonal direction. **d.** Estimation of the experimental localization uncertainly projected on the surface of the MT. **e,** Estimated apparent energy landscape considering the experimental uncertainly. **f,** Longitudinal and transverse cross-sections of the estimated apparent energy landscape along the dashed lines indicated in e. Scale bars in b, d, e equal 4 nm.

The set of simulated trajectories of the Ase1 in the plane projected to the MT surface overlaid with the resulting density map of positions summed in one tubulin-dimer cell and extended periodically for better visualization to a 3×3 patch of the lattice of tubulin dimers is shown in Figure 5b. The simulated trajectory of Ase1 clearly shows transient confinements localized with different probabilities at the two dominant sites of the tubulin dimer. During the simulation, occasional transitions between tubulin dimers can be observed. In particular, we identified 13 longitudinal passes along the MT axis, five transversal jumps between the protofilaments, and three more complex diagonal transitions (Figure 5c). Despite the low statistical significance of the transition events between tubulin dimers, it is interesting to note that their occurrence rate closely agrees with the experimental stepping probability discussed in the previous section (Figure 4e).

Molecular dynamics (MD) simulations traditionally serve as a virtual in silico microscope that can reach the atomic level and allows a tentative interpretation of the experimental results. Unfortunately, these experiments usually have no comparable resolution. Encouraged by the close agreement of the theoretical model with the observed behavior of the Ase1, we scrutinized the extraction of the Ase1-tubulin interaction potential with the experimental localization uncertainty. Therefore, we characterized the localization uncertainty of our experimental trajectories (Supplementary Fig. S2); the estimated uncertainty distributions projected at the surface of the MT with a 2.1-nm and 1.1-nm standard deviation in the longitudinal and the transversal direction, respectively, are depicted in Figure 5d. We estimated the probability of localizing the Ase1 molecule at different positions of the MT lattice and derived the corresponding potential energy profile *U*(*x,y’*) (Figures 5e and 5f). Although fine details of the interaction potential on the level of one tubulin dimer were lost in the overall energetic landscape, our estimation indicated that the strength of individual Ase1 confinements to a single-tubulin dimer should be distinctly resolvable experimentally.

### Experimental characterization of the interaction potential

One of the selected Ase1 trajectories (fast diffusivity of *D_x_*=65 nm^2^ ms^-1^, Figure 3d), binned to a 335-μs temporal resolution, corrected for the MT supertwist geometry, and split into regions of 24-nm (three tubulin dimer) period is shown in Figure 6a. These regions were superimposed and accumulated to obtain the Ase1 position histogram on the three-tubulin-dimer long segment of MT (Figure 6b). The same accumulation procedure was performed with an 8-nm (one tubulin dimer) long region and a sequence of three identical 8-nm segments of the accumulated position histogram is shown in Figure 6c to emphasize the periodicity of the resulting pattern visually matching the 24-nm segment in Figure 6b (Methods). The position of the five horizontal bands of increased probability density closely agreed with the expected spacing of individual protofilaments of 6 nm. The profile features a pattern of locally increased probabilities, highlighted with red circles in Figure 6c. Interestingly, this pattern features a shear between neighboring protofilaments of approximately 1.3 nm which is closely matching the pattern of tubulin dimers obtained from cryo-EM studies of the MT lattice. Furthermore, the pattern of the apparent position of the tubulin dimers revealed a 4-nm longitudinal offset in the periodicity between the lower second and third protofilament (highlighted using red circles in Figure 6c). This geometry closely matches the arrangement of tubulin dimers in the so-called seam of the MT.^39^

**Figure 6.**
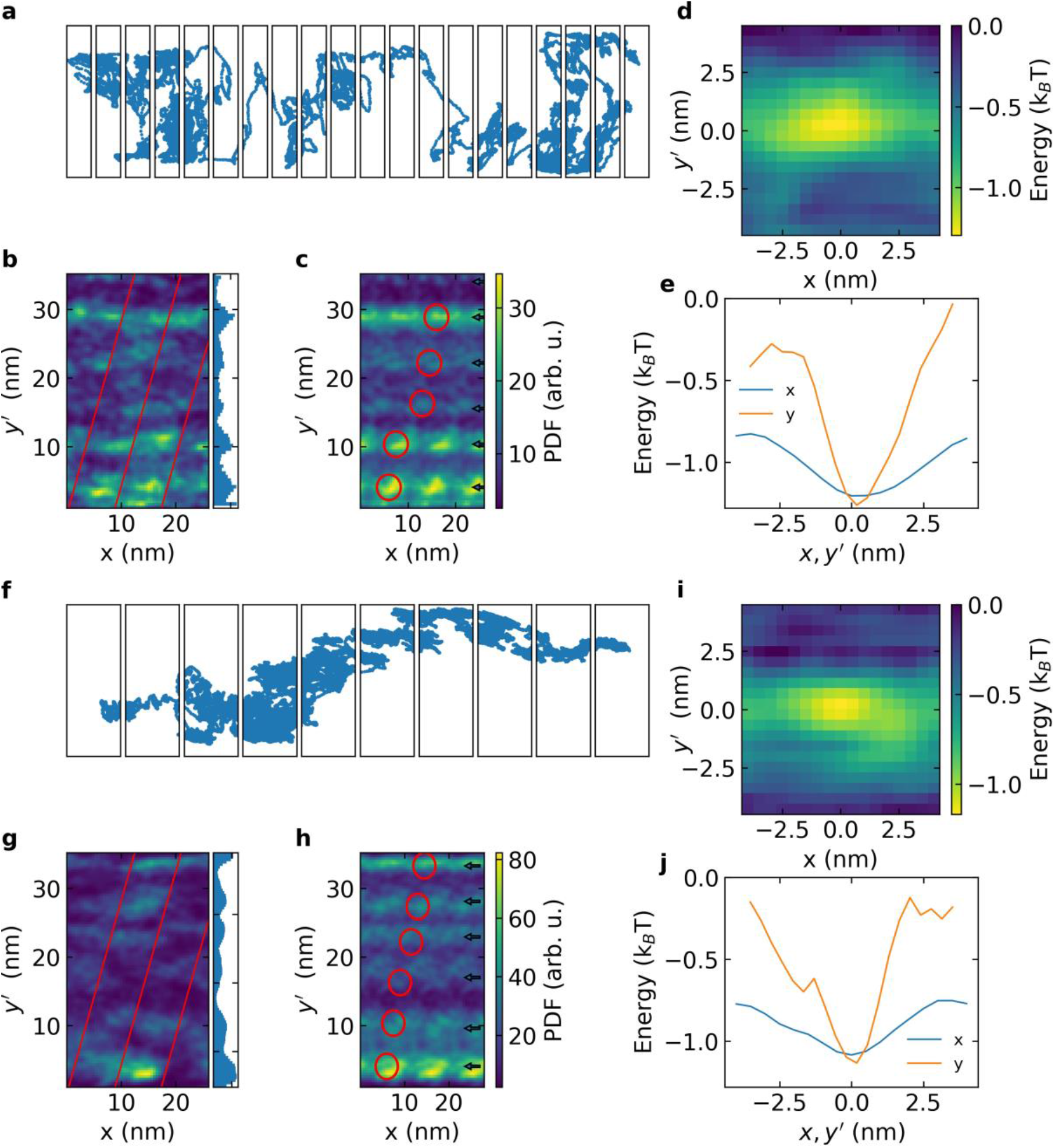
Extracting the single-tubulin-dimer interaction potential. **a, f,** Two examples of Ase1 trajectories (smoothed) split into sections of 24 nm (three tubulin dimers) along the MT axis. **b, g,** 2D histograms of the localized position in the stack of 24-nm sections indicated in a, f. **c, h,** 2D histograms of the localized position stack of 8-nm sections (three identical periods shown along x). **d, i,** The mean 2D interaction potential of the Ase1 with one tubulin dimer. **e, j,** Longitudinal and transverse cross-sections of the measured interaction potentials.

The same analysis was performed for another example trajectory showing diffusivity of *D_x_*=3.1 nm^2^ ms^-1^, which corresponds to the slow-end of the experimental span in Figure 6f-h. The resulting localization density map in figure 6h features a similar pattern corresponding to the 6-nm spacing between individual protofilaments as well as the increased probability of Ase1 localization at the tubulin-dimer periodicity.

We averaged the mean profile of the probability density function *p*(*x,y’*) from one tubulin-dimer proximity of all five equilibrium positions of the Ase1 in the accumulated position histogram in Figure 6c,h and used it to reconstruct the potential energy profile *U*(*x,y’*) using the equipartition theorem. Figures 6d and 6i represent the resulting potential wells, indicating an energetic barrier of approximately 1 k_B_T in the transversal direction, whereas the energy barrier along the MT was found to be approximately 0.4 k_B_T. Therefore, the extracted landscape of the confinement closely agreed with the theoretical model in Figure 5. We note that the fast fluctuation of the GNP label limits the details of the energetic landscape extraction at the nanometer level apparent from the comparison of Figures 5b and both 5e,j. The maximum resolvable gradient of the interaction potential reached approximately 0.1 k_B_T nm^-1^ in the longitudinal direction and 0.2 k_B_T nm^-1^ in the transversal direction (higher owing to the scaling effect between the position of the GNP and the surface of the MT). Therefore, the maximum gradients of the interaction potentials resolved in Figures 6e and 6j are very close to the measurable limit, and estimation of the depth of the potential well shows only a lower bound assessment. It is worth noting, that the resulting profile in Figure 6i-j yielded an almost identical potential well compared with that in Figure 6d-e, even though all other measures including the dwell time statistics, the displacement histogram and the overall diffusivity indicated stronger interaction between the Ase1 and MT. We attributed the close match of the two extracted profiles of the potential wells to the quantitative limitation of the maximum interaction potential gradient assessment. Despite the experimental limitation in estimating the full depth of the potential well at an atomic resolution, the outstanding spatial and temporal resolution of our data enabled a very close match between the MD simulation and the high-fidelity tracking experiment, providing a unique validation of the complex theoretical model.

## Discussion

Diffusion is one of the essential strategies that proteins use to navigate the complex environment of the MT surface. It is employed by a large variety of MAPs, including molecular motors,^1,40-42^ intrinsically disordered neuronal proteins,^9,43–45^ proteins tracking polymerizing or depolymerizing microtubule tips^4,9,46,47^ or microtubule crosslinker proteins.^4,9,15,48^ Stepping of proteins from the Ase1/PRC1/MAP65 family across protofilaments has been predicted theoretically^9^, and here we present a direct experimental validation of this prediction. The ability to step across protofilaments has been shown to enable molecular motors to navigate around obstacles on the microtubule lattice.^49,50^ We deciphered the mechanism underlying the diffusive interaction of MT crosslinker Ase1 with the MT surface by analyzing the complex trajectory of Ase1 on a single MT at a very high spatial and temporal resolution. We discerned nanoscopic confinements in the diffusive trajectories and associated them with individual equilibrium positions of Ase1 interacting with tubulin dimers within the MT lattice. Step detection analysis indicated a mean dwell time of these confinements of 1.6 ms followed by a displacement of the molecule to the adjacent tubulin dimer, with the step size of a single tubulin dimer being the most probable, in accordance with previous experimental^4^ and theoretical^9^ work. Interestingly, our analysis revealed an approximately two-fold higher probability of Ase1 stepping in the direction of the MT axis compared with the transversal displacements in the perpendicular direction. This conclusion was consistent with the overall diffusivity analysis, which yielded a similar ratio of the longitudinal and transversal diffusivity at the time-scales exceeding the confinement dwell time. Our results showing the probability distribution of Ase1 stepping indicate that Ase1 can explore the MT length while efficiently navigating obstacles. This feature might be of importance, especially in the highly crowded MT overlaps that Ase1 stabilizes.

Considering the diffusion of Ase1 in strictly periodic boundaries, we extracted the landscape of the interaction potential in the vicinity of the equilibrium confinements. Indeed, we were able to reconstruct the shape of a potential well associated with the interaction with a single tubulin dimer and independently quantify the energetic barrier in the longitudinal and transversal direction. In agreement with the theoretical prediction, we showed that Ase1 experiences a larger energetic barrier when jumping between protofilaments compared with the displacement along a single protofilament. We observed two effects influencing the final long-range diffusivity of the protein on the MT surface: the higher the diffusivity of the Ase1 protein (i) the shorter the mean dwell time in each confinement of the trajectory and at the same time (ii) the larger displacement between individual confinements can be detected. Combined, the effect of the dwell time variation together with the displacement between confinements results in the variation of the measured diffusivity exceeding one order of magnitude.

Diffusive transportation along microtubules is employed by a large variety of proteins. Using iSCAT microscopy with a high temporal and 3D spatial resolution, we imaged the diffusion of an exemplary diffusible MAP over a microtubule lattice, which enabled us to resolve the preference of Ase1 to move along the protofilament in single tubulin-dimer steps. Experimental characterization of interactions at the single-protein level with very high accuracy and precise details at the same time, allowed us to close the gap between *in silico* and optical microscopy. We hypothesize that stepping characteristics underpin various functional properties of MAPs. Therefore, it would be intriguing to determine the stepping characteristics of other diffusible MAPs, including diffusible motor proteins, and obtain new insights into the dynamics of single-protein machinery.

## Materials and Methods

### Materials

All chemicals were purchased from Sigma Aldrich (Merck) unless stated otherwise and used as received without further purification. Ultrapure water from a Direct-Q 3 UV water purification system (Millipore, USA) was used for all experiments.

### Flow chamber preparation

18×18 mm (top) and 22×22 mm (bottom) coverslips were rinsed with ethanol and water. After drying with N_2_ both coverslips were cleaned with oxygen plasma (Diener, Germany) to remove possible impurities from the surface. Clean bottom coverslips were silanized as follows: First, they were dipped in acetone for 10 s and then immersed in a 2 wt. % solution of 3-aminopropyl triethoxysilane (APTES) in acetone for another 10 s. Finally, the bottom coverslip was rinsed with acetone and dried with N_2_. The top and bottom coverslips were assembled with two parallel spacers of Parafilm M. Each assembled coverslip was heated to seal the flow chamber.

### Ni-NTA functionalization of gold nanoparticles

Functionalization of the gold nanoparticles (GNPs) with the nickel-nitrilotriacetic (Ni-NTA) functional group was performed because Ni-NTA selectively binds to the His-tag region of the protein.

The surface of the citrate-stabilized GNPs (BBI solutions, United Kingdom) was modified with NTA-terminated thiols (3,3’-dithiobis[N-(5-amino-5-carboxypentyl)propionamide-N,N’-diacetic acid] dihydrochloride) (dithiobis(C2-NTA)), Dojindo, Japan), followed by the addition of a Ni^2+^ salt and its complexation with the NTA group.

In the first step, 300 μL of 40-nm citrate-stabilized GNPs were cleaned by centrifugation at 9000 rpm for 10 min and resuspended in 200 μL of 20 mM (4-(2-hydroxyethyl)-1-piperazineethanesulfonic acid) (HEPES) buffer with 0.1% Tween 20. Then, these pre-cleaned GNPs were mixed with 100 μL of a 1.3 mM dithiobis (C2-NTA) solution and left overnight. The thiol molecules chemisorb on the gold and, thus, cover the entire nanoparticle surface. The unreacted thiol was removed by several centrifugation cycles at 7000 rpm for 6 min and resuspended in 200 μL of 20 mM HEPES buffer with 0.1% Tween 20.

Next, 1 μL of a 10 mM NiCl_2_ solution was added to 100 μL of the NTA-functionalized GNPs and left for 10 min at room temperature. Finally, GNPs were washed by centrifugation and resuspended in the same amount of HEPES 0.1% Tween 20. A schematic representation of the procedure is shown in Supplementary Fig. S8.

### Double-stabilized MT synthesis

Porcine tubulin was purified from pig brain according to Castoldi and Popov.^51^ To polymerize MTs,^42^ the purified tubulin was incubated for 2 h at 37 °C in a 100 μL solution containing BRB80 (80 mM piperazine-N,N′-bis(2-ethanesulfonic acid) (PIPES) pH 6.9, 1 mM MgCl_2_, 1 mM ethylene glycol-bis(2-aminoethyl ether)-*N,N,N′,N′*-tetraacetic acid (EGTA)), 1 mM GMPCPP (Jena Bioscience, Germany), 1 mM MgCl_2_ and 4 mg mL^-1^ of porcine brain tubulin. Assembled MTs were centrifuged at 12000 g for 30 min. The pellet was resuspended in 100 μL BRB80/Tx (BRB80 containing 10 μM of Taxol).

### Interferometric scattering microscopy (iSCAT) setup

The output of a continuous wave laser emitting at 561 nm (Laser Quantum, United Kingdom) with a power of 16 mW cm^-2^, was circularly polarized, modulated with an acousto-optic modulator (AA Opto Electronic, France), synchronoized with the camera frame exposure, and spatially filtered into a speckle-free beam. The illuminating beam was then transmitted through a 70:30 beam splitter and propagated through the focus (at the back focal plane) of a high-NA microscope objective (α Plan-Apochromat 100x, 1.46 NA, Carl Zeiss AG, Germany) to illuminate an area of 4.5 μm × 4.5 μm on the functionalized coverslip. Light reflected on the glass-water interface and light scattered by a nanoparticle in the vicinity of the glass surface was collected by the same microscope objective and redirected at the 70:30 beam splitter to be imaged on a CMOS-based camera (Photonfocus AG, Switzerland).

The flow chamber containing the liquid sample was mounted on top of a 3-axis piezo stage (Physik Instrumente GmbH & Co. KG, Germany), to position the sample with sub-nanometer precision.

### Fast-tracking of single Ase1

Prior to the experiment, the flow chamber was mounted to the microscope head. Then, this flow chamber was filled with BRB/Tx. Next, 10 μL of double-stabilized MTs were injected. The MTs exhibit a negative net charge due to the low isoelectric point of the tubulin subunits below 6.0. As a result, they were electrostatically immobilized on the positively-charged APTES surface. The remaining free APTES was passivated with a 10 mg mL^-1^ β-casein solution for 2 min.

Afterwards, the flow chamber was flushed with the motility buffer containing 20 mM of HEPES, 2 mM of MgCl_2_, 1 mM EGTA, 75 mM KCl, 10 μM taxol, 0.5 mg mL^-1^ casein, 10 mM dithiothreitol (DTT), 1 mM ATP(+Mg), 20 mM D-glucose, 220 μg mL^-1^ glucose oxidase and 20 μg mL^-1^ catalase. The Ase1 protein, diluted with the motility buffer to 0.60 nM concentration, was injected into the flow chamber. We ensured that the Ase1 concentration is low enough, such that the mean Ase1 to Ase1 distance on the microtubule was 3±1 μm, enabling a single-molecule study (see Supplementary Information for a control experiment using single-molecule fluorescence). After 5 minutes, 5 μL of the Ni-NTA GNPs were injected and incubated for 2 min. Finally, the GNP solution was exchanged for the motility buffer.

The strong affinity and selectivity of the Ni-NTA linker to the His-tag of the Ase1 molecule enabled fast labeling (incubation time of 2 min) of Ase1 molecules diffusing along the MT surface, as verified by comparing the final labeling density with a single-molecule fluorescence experiment (Supplementary Information). Within the time window until the majority of nanoparticles are attached to the Ase1, an average Ase1 molecule displaces less than 2.5 μm along the MT (D~55 nm^2^ ms^-1^)^15^, which is less than the mean Ase1-to-Ase1 distance on the MT.

### Image acquisition and data processing

A custom-written LabView software (National Instruments) was used to control the experiment and collect data. For a real-time preview of the illuminated area of 128 × 128 pixels (~4.5 × 4.5 μm) during the experiment, a framerate of 7000 frames per second was used.

Upon identifying a single diffusing Ase1 on the MT, the exposure time was set to 10 μs, and high-speed image data within a region of 32 × 32 pixels (~1250 × 1250 nm) were acquired at a 45 kHz frame rate. To suppress the effect of the static background, a reference measurement (1000 frames) was acquired immediately after each fast measurement. A series of reference images were recorded at a position displaced by a few micrometers within an area showing no strongly scattering features using the same settings as for the tracking measurement. Reference images were averaged and subtracted from the high-speed image data.

Particle tracking was performed using the open-source TrackMate plugin for ImageJ^52–54^ to determine both x- and y-positions as well as the contrast for each recorded frame. For visualizing and data processing, we used the Python 3.6 programming language (Python Software Foundation, https://www.python.org/). XY coordinates were transformed to align the MT axis along the X coordinate.

Mean square displacement (MSD) analysis was performed on the XY single-particle trajectories. At given *Δt_n_* time delay the 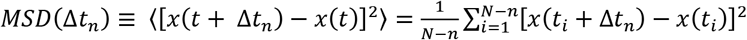, in which N is the number of localizations in the trajectory, *n* = 1,…, *N* - 1 is the number of localizations at *Δt_n_* time delay.

Diffusion constants *D_x_* were obtained by fitting of the MSD curve with the equation: *MSD*(*Δt*) = ϵ + *2nDΔt*, in which ϵ is the localization uncertainty, n=1 is the number of free dimensions of the diffusive motion, and *Δt* is the time lag between positions. ^55,56^

To estimate the diffusivity *D_y′_* we calculated the mean ratio 〈*MSD_y_/MSD_x_*〉 of the transverse and the longitudinal MSD in the range from 1 to 3 ms. The transverse diffusivity coefficient was estimated as *D_y_* = *D_x_*〈*MSD_y_/MSD_x_*〉.

### Extraction of the MT interaction potential

To extract the profile of a single tubulin dimer interaction potential, we first rotated the grid of x-y’ data to ensures that the protofilaments of MT were parallel to the x-axis, accounting for the supertwist angle and orientation. Then the trajectory was split along the x-axis at 24-nm or 8-nm intervals and 2D histograms of localizations within the sum of the split regions were calculated. Variation of the 24-nm and 8-nm splitting interval was tested to obtain the maximum contrast of the 2D histogram of localizations.

### Indexing the steps between the trajectory confinements

Trajectories were smoothed with a 335-μs time window and a 2D histogram of the trajectory was calculated with a 0.4-nm discretization. The histogram was smoothed to a spatial resolution of 1.6 nm using a Gaussian filter. All local maxima were detected in the resulting 2D histogram and the localizations within their surrounding within the decreasing gradient were indexed as belonging to each selected local maximum. Regions corresponding to the vicinity of each local maximum are incrementally color-coded in Figure 4 and Supplementary Fig. S6.

Dwell times were extracted as resident time in the vicinity of each local maximum. Histograms of dwell times were calculated with an exponential discretization and fitted to an exponential decay from 1 ms to 1 s using a weight function proportional to the bin size. The displacement vector was calculated from the mean positions of respective local maxima at every detected step. The 2D histogram of all displacement vectors represents the probability map of steps for each trajectory separately.

### Coarse-grained molecular dynamic simulations

Recently, coarse-grained MD simulations have been successfully used to study protein diffusion on microtubules and DNA.^9,16^ We followed the methodology from these papers. A coarse-grained Cα-based model of an MT lattice and diffusing Ase1 protein was generated using the SMOG web server (www.smog-server.org^57,58^). The applied force-field used a model based on the native topology.^57,58^ Native contacts were rewarded by means of a Lennard-Jones potential. Non-native contacts were penalized using a repulsive potential. Electrostatic interactions between charged residues were modeled using the Debye-Huckel potential.^59^ The flexibility of the disordered C-terminal tails of tubulins as well as Ase1 was controlled by its angle and dihedral angles force constants. The dynamics of Ase1 diffusion along MT was simulated using the Langevin equation using GROMACS 5.0.5 ^60^. The MD simulation temperature was set to 50 (in reduced units), which is lower than the folding temperature of proteins (100-120) in coarse-grained MD simulations. The dielectric constant was 80 and the salt concentration was 0.03 M. MD simulations consisting of 3.7×10^9^ essential MD steps (0.0005 in reduced units) were produced. Trajectory frames were saved every 20000 steps. Simulated systems were visualized using the VMD 1.9.4 software package^61^ and rendered using Blender 2.91.0 (www.blender.org).

## Data availability

Data supporting the findings of this study are available in the Supplementary Information and all raw data are available from the corresponding author upon reasonable request

## Acknowledgments

This work was funded by the Ministry of Education, Youth and Sports of the Czech Republic under project LL1602, the Czech Science Foundation under project 19-27477X, and the institutional support from the CAS (RVO: 86652036).

## Author Contributions

K.H. and Ł.B. developed the optical setup, K.H, Ł.B., and A.G.M. performed the experiments; Ł.B. and M.P. processed and interpreted the data; I.B. performed MD simulations, V.H. generated the proteins, M.P., Z.L. and M.B. conceived the research; Ł.B., M.P., and Z.L. interpreted the results and co-wrote the paper.

