## Supplementary Information for "Fast leaps between millisecond confinements govern Ase1 diffusion along microtubules"

### 1. Instrumental localization precision

We characterized the localization precision of a 40-nm static nanoparticle on the glass surface. Negatively-charged citrate-stabilized GNP was immobilized on a positively-charged APTES-coated surface to characterize the localization precision achievable with the label. The position of the GNP was recorded and further localized with ImageJ. Results are shown in Figure S1 for different planes. The standard deviation of the Gaussian fit in x-, y- and z-axis was $\sigma_x$=0.55 nm, $\sigma_y$=0.83 nm, $\sigma_z$=0.62 nm, respectively.

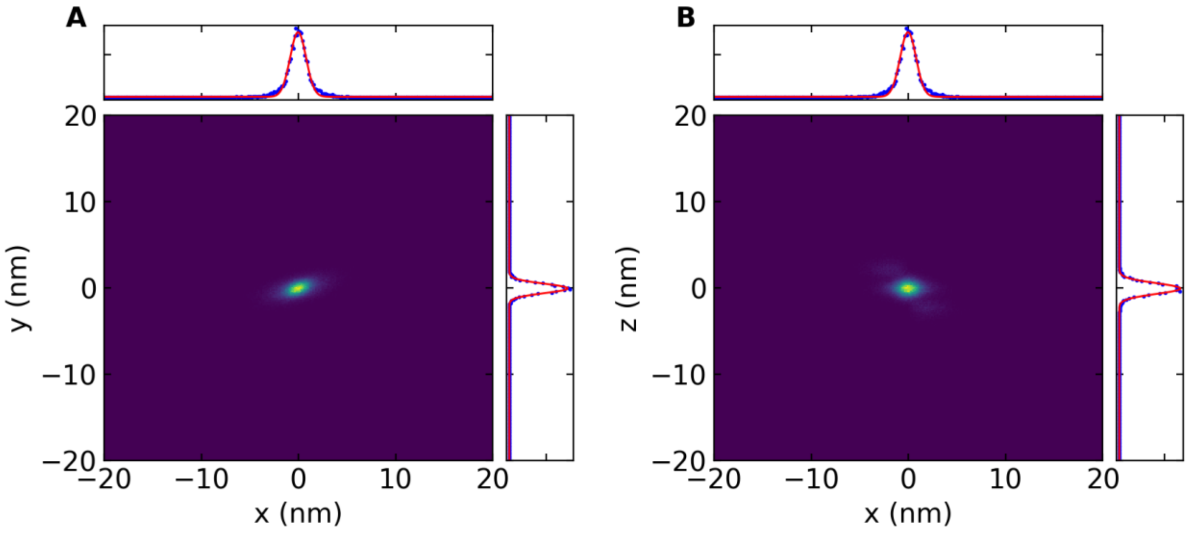


Figure S1. A Probability histogram of the apparent positions of an immobilized GNP in (A) x-y plane and (B) x-z plane.

### 2. Localization precision of a diffusing MAP

Next, we analyzed the noise component of the trajectory of the Ase1-GNP construct. We extracted the single-frame 3D displacement through all the experiments and Figure S2 shows the histogram of the localization. The statistics of the localizations suggests Gaussian distribution of rather uniform standard deviation $\sigma_x$=3 nm, $\sigma_y$=3.1 nm, $\sigma_z$=3 nm. The single-frame particle displacement is, thus, not limited by the localization precision of the GNP itself. Instead, it likely reflects the real diffusive motion of the GNP attached to the Ase1 protein. Interestingly, the statistics of the 3D localization is close to a spherically symmetrical pattern. This spherically symmetric behavior is probably associated with relatively loose attachment of the GNP to the MT via a flexible anchor mediated by the Ase1 molecule. The 3-nm diameter of the histogram of the single-frame GNP displacement can be interpreted as a convolution of two consecutive random fluctuations in the XYZ position. Following this assumption, we estimate that the precision of the Ase1 localization based on the GNP position is approximately 2.1 nm.

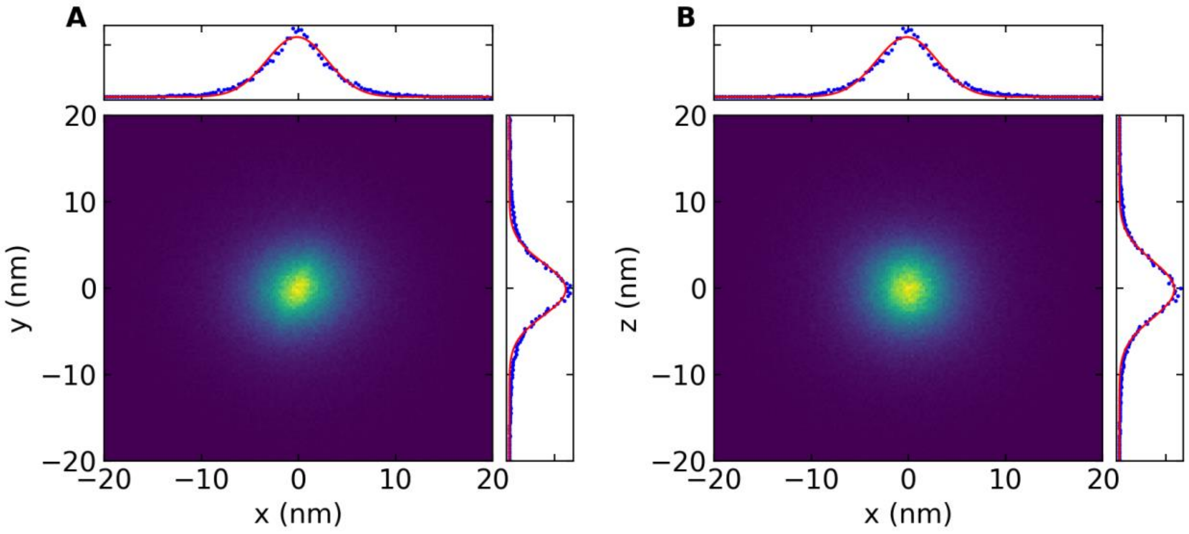


Figure S2. Probability histograms of the single-frame increment in the apparent position of Ase1-GNP construct on the MT in (A) x-y plane and (B) x-z plane.

#### 3. Estimation of Ase1 density using single-molecule fluorescence detection

In order to ensure the single-molecule regime in our experiments, we performed a control using single-molecule fluorescence detection. To do that, we use a green fluorescent protein (GFP) labeled mutant of the Ase1 molecule and coupled the fluorescence beam path with the iSCAT microscope using a dichroic mirror (FF495-Di03-25x36, Semrock, USA) and dichroic filter (FF02-520/28-25, Semrock, USA) with the EMCCD camera (iXon, Andor, UK). For excitation, we used a 445 nm laser diode (Lasertack, Germany).

The experimental procedure described in the Materials and Methods section was the same for the fluorescence control experiment. In this case, we injected the Ase1, labeled with the GFP as well as with a His-tag available in the dimerization region, with concentrations ranged between 5.40 and 0.45 nM. Figure S3 shows four examples corresponding to different Ase1-GFP concentrations incubated with the previously-attached MTs on the glass channel. For the concentration of 5.40 nM, the MTs were covered with the fluorescent proteins at a high density not allowing to resolve single proteins. With concentration decreasing to the nanomolar and sub-nanomolar range, single fluorescent proteins were resolved to diffuse along the MT. To estimate the protein density at the conditions used for the tracking experiments, we used the Ase1-GFP concentration of 0.60 nM. We scanned an area of 930 $\mu m^2$ in the fluorescence video, where we found a total number of Ase1-GFP of 54, yielding a value of 0.06 Ase1-GFP $um^{-2}$. Similarly, we examined an area of 300 $\mu m^2$ in the iSCAT video where we found an MT total length of 59 µm, yielding an average MT density of 0.2 µm $\mu m^{-2}$. With both values, we estimated a protein density of 0.30 Ase1-GFP $\mu m^{-1}$ of MT (i.e. ≈ 1 Ase1-GFP per 3 µm of MT).

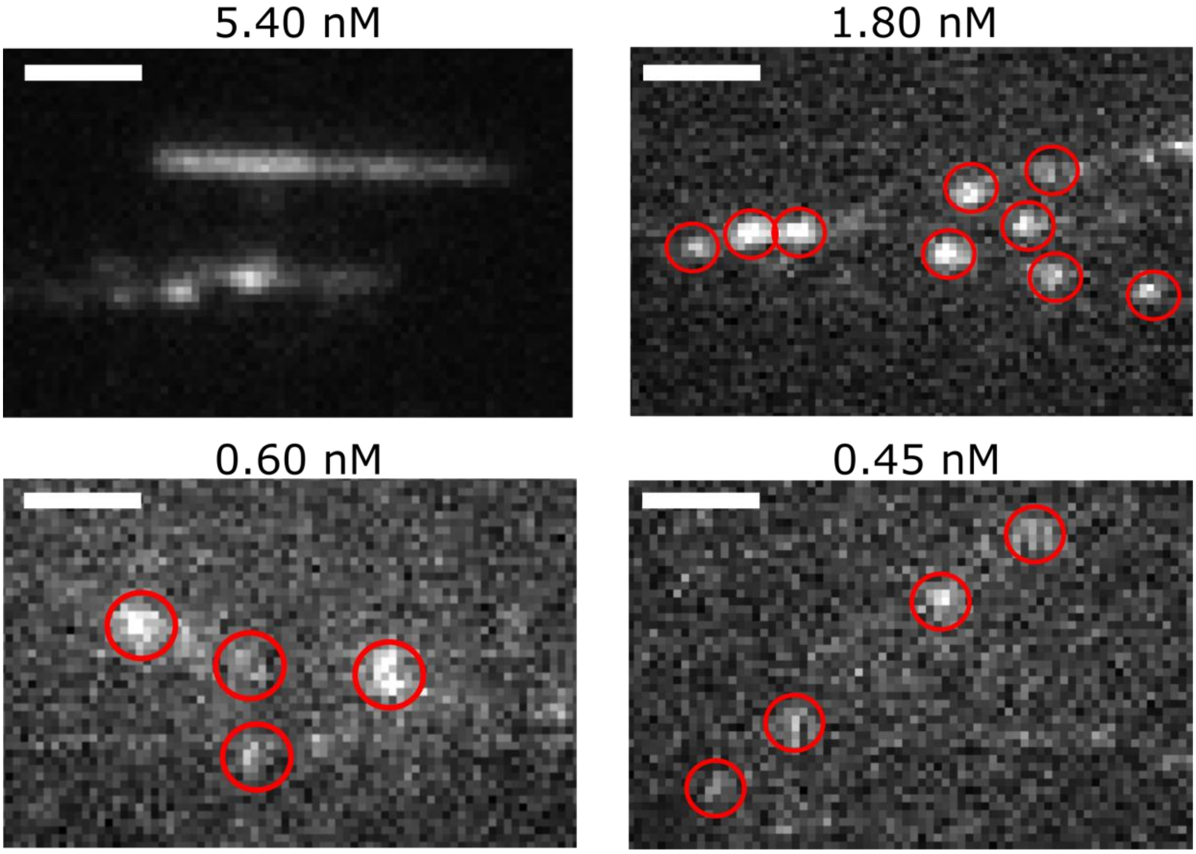


Figure S3. Wide-field fluorescence microscopy images of the dilution series of Ase1-GFP on MT. The protein concentrations incubated with the microtubules inside each sample are noted. Scale bars represent 1.0 µm.

**4. GNP – Ase1 coupling efficiency**

To explore the efficiency of the GNP attachment to Ase1 we recorded the scattering and fluorescence signal simultaneously in the same field of view. It is worth noting, that this experiment requires significantly lower illumination power (0.5 mW $cm^{-2}$) to allow for sufficient observation times of the fluorescence. Therefore, the iSCAT control in this experiment is considerably noisier and this implementation is therefore used solely in this control experiment. We used the same protocol described in Methods (Fast-tracking of single Ase1). The Ase1-GFP at a concentration of 0.60 nM was injected and incubated for 5 minutes with MTs. Next, the 0.15 nM solution of 40-nm Ni-NTA GNPs was introduced for two minutes to label the protein with the GNPs. Figure S4 shows an example of the simultaneous iSCAT and fluorescence images recorded at the same position. The Ase1-GFP proteins labeled with the GNPs are highlighted with red circles. A clear match between the fluorescence signal and the detected position of the GNP labels already indicates high labeling efficiency. We estimated an average GNP labeling efficiency of 50 ± 20% based on the statistical analysis of 300 $\mu m^2$ of the surface.

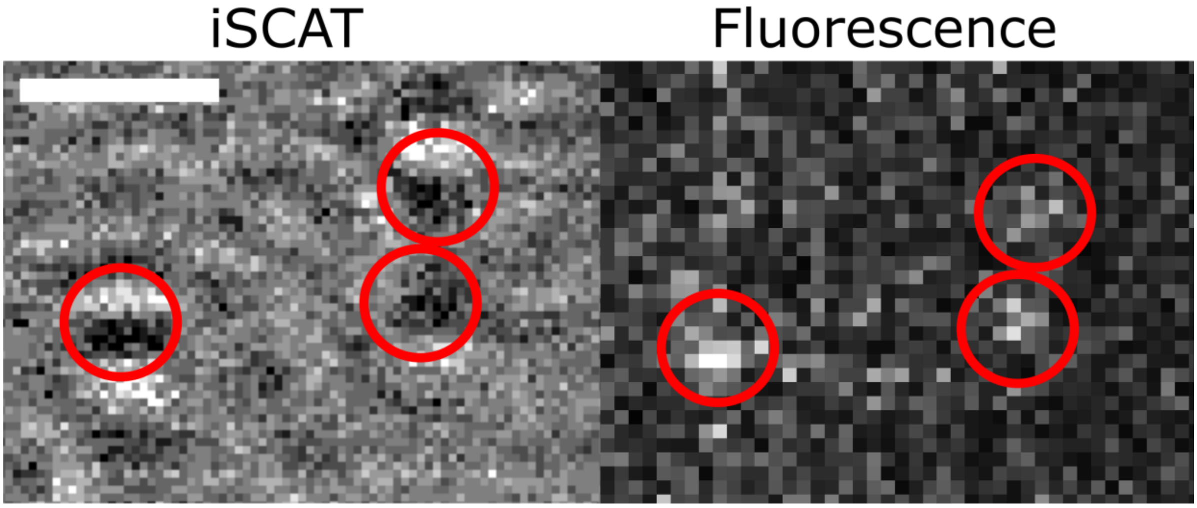

Figure S4. Comparison of iSCAT and fluorescent images of Ase1 (0.60 nM) labeled with the GFP protein and 40-nm Ni-NTA GNP. Red circles indicate GFP and GNP detected signals. Scale bar represents 1.0 µm.

#### 5. Directional probability of the Ase1 stepping

Probabilities of displacements to a specific tubulin neighbor in Supplementary Figure S7 were calculated from the two-dimensional histograms of the displacement vector

$$\mathrm{p}_i = \frac{\sum_{j,k=0}^{N_i} d_{j,k}^i}{\sum_{j,k=0}^{N} d_{j,k}} * 100$$

Where $\mathrm{p}_i$ is probability of displacement from the center to the $i$-th grid cell, $d_{j,k}^i$ is number of occurrences at positions $j, k$ within each $i$-th grid cell, $N_i$ number of all positions in $i$-th grid cell, $d_{j,k}$ is the number of occurrences at positions $j, k$, $N$ number of all positions in displacements histogram. The resulting probabilities $\mathrm{p}_i$ are indicated in Figure S6 in each corresponding grid. To calculate probability of stepping along MT we summed all probabilities of displacement along MT ($p_x$) and divided it with the sum of probabilities along the central protofilament and perpendicular to it ($p_y$).

$$p_x = \frac{\sum_{i=0}^{N} p_x^i}{\sum_{i=0}^{N} p_x^i + \sum_{i=0}^{N} p_y^i}$$

$$p_y = \frac{\sum_{i=0}^{N} p_y^i}{\sum_{i=0}^{N} p_x^i + \sum_{i=0}^{N} p_y^i}$$

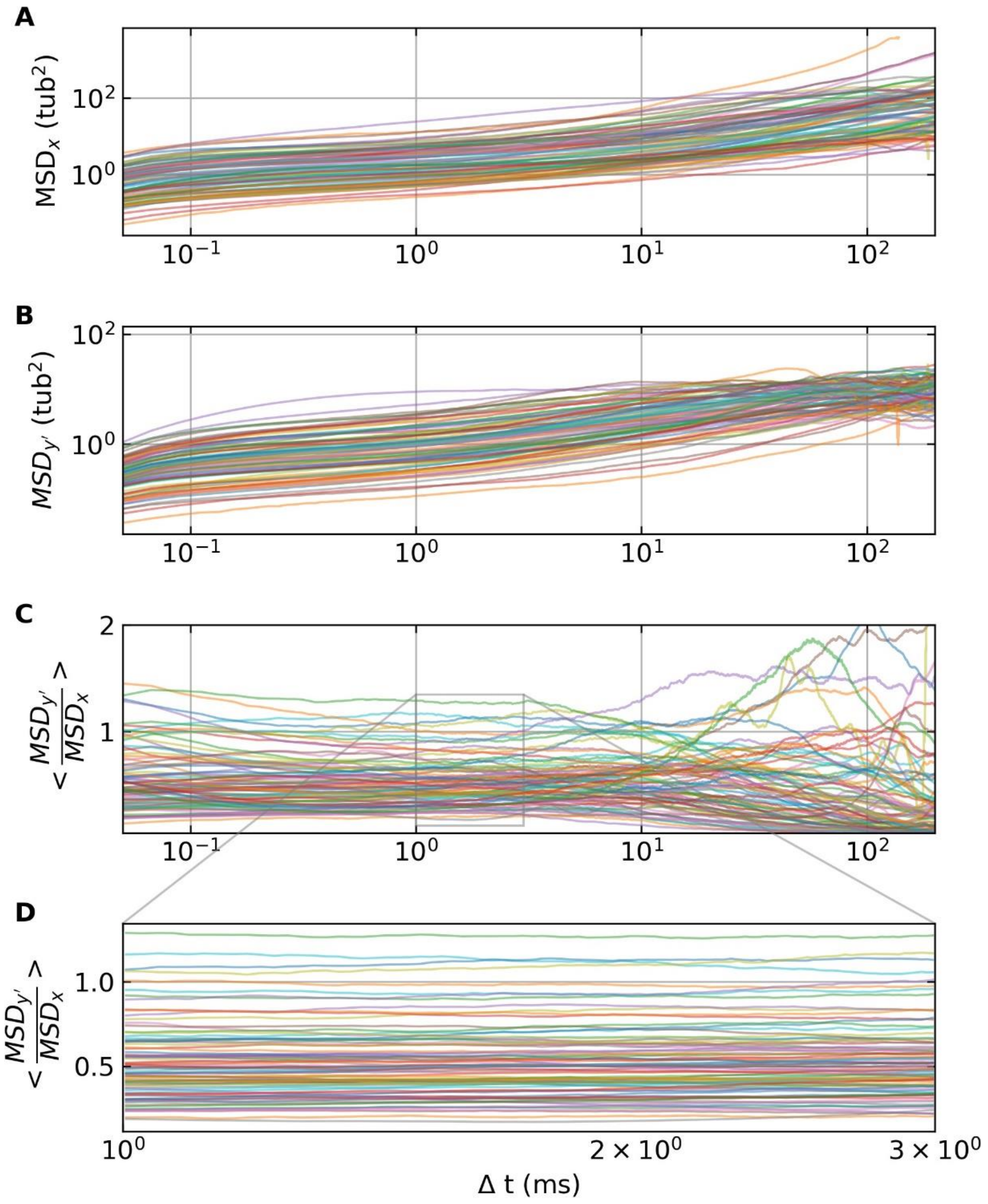


Figure S5. Comparison of MSD plots calculated for all measured traces. (A) MSD curves calculated from the longitudinal displacement; (B) MSD curves calculated from transversal displacement; (C) ratio of the transversal and longitudinal MSD curve and (D) zoom into the linear segment of this ratio.

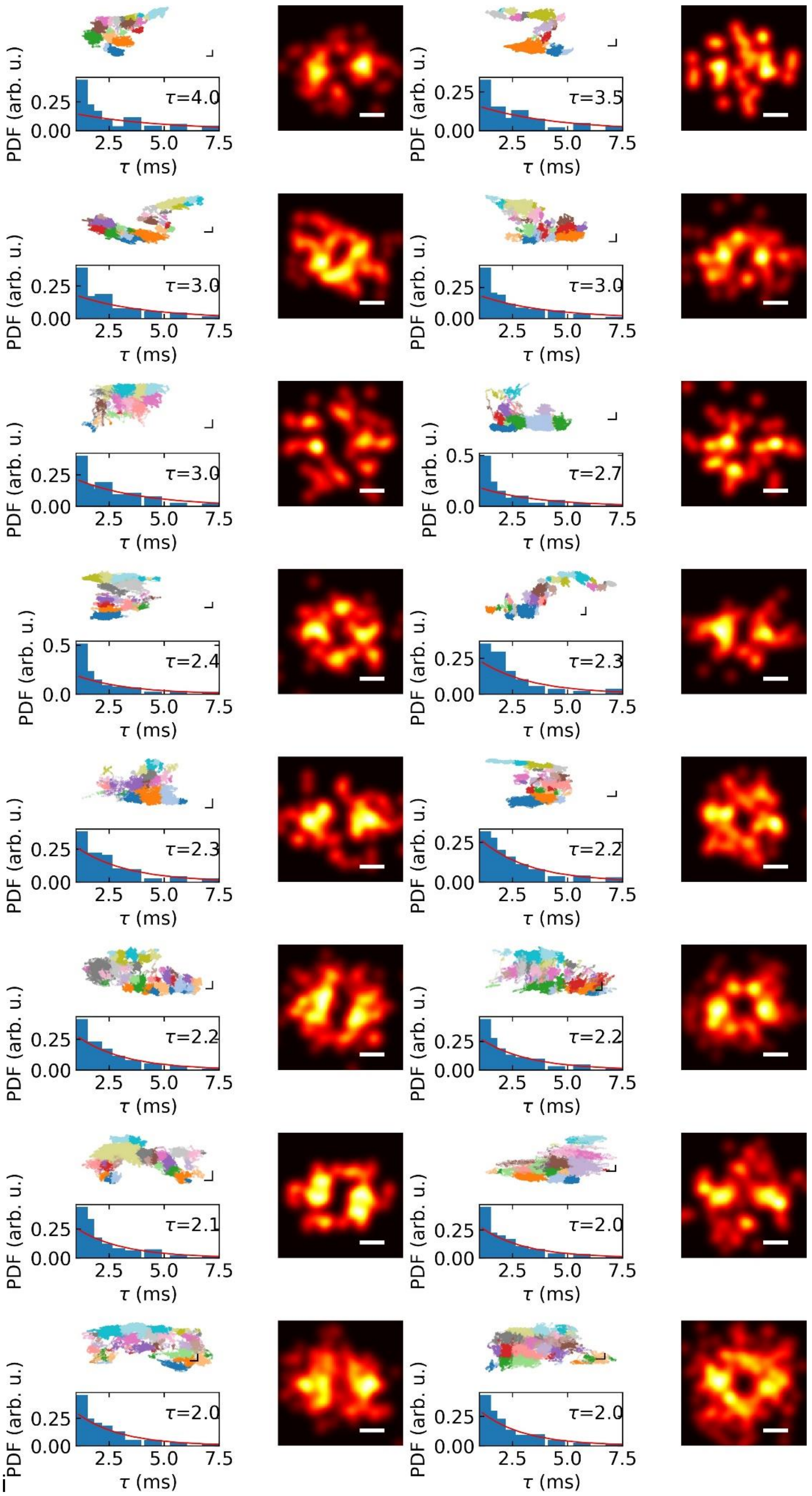

PDF (arb. u.)
τ (ms)
τ=4.0
τ=3.5
τ=3.0
τ=3.0
τ=3.0
τ=2.7
τ=2.4
τ=2.3
τ=2.3
τ=2.2
τ=2.2
τ=2.2
τ=2.1
τ=2.0
τ=2.0
τ=2.0

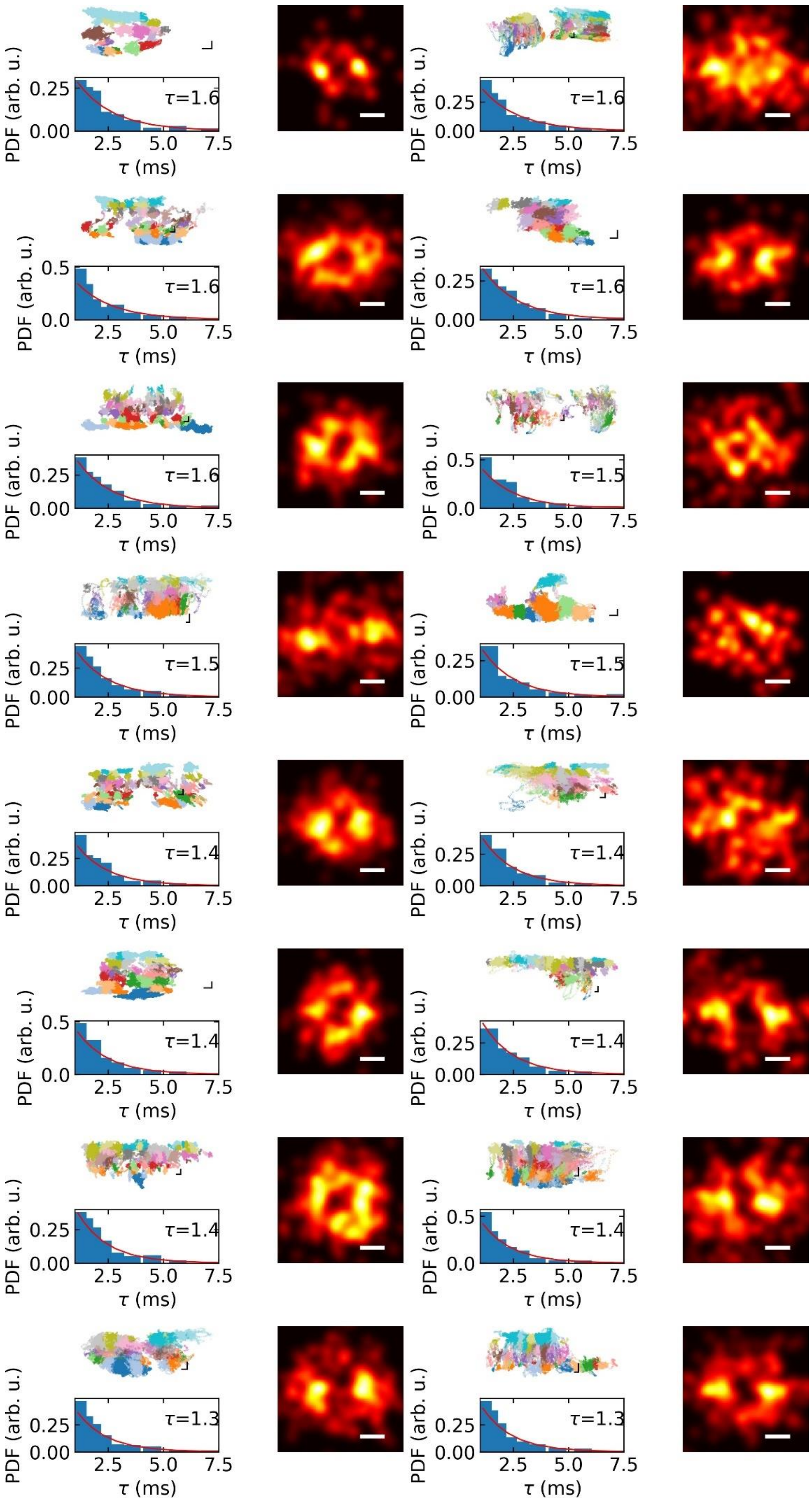

PDF (arb. u.)
0.25
0.00
τ=1.6
2.5 5.0 7.5
τ (ms)
PDF (arb. u.)
0.25
0.00
τ=1.6
2.5 5.0 7.5
τ (ms)
PDF (arb. u.)
0.5
0.0
τ=1.6
2.5 5.0 7.5
τ (ms)
PDF (arb. u.)
0.25
0.00
τ=1.6
2.5 5.0 7.5
τ (ms)
PDF (arb. u.)
0.25
0.00
τ=1.6
2.5 5.0 7.5
τ (ms)
PDF (arb. u.)
0.5
0.0
τ=1.5
2.5 5.0 7.5
τ (ms)
PDF (arb. u.)
0.25
0.00
τ=1.5
2.5 5.0 7.5
τ (ms)
PDF (arb. u.)
0.25
0.00
τ=1.5
2.5 5.0 7.5
τ (ms)
PDF (arb. u.)
0.25
0.00
τ=1.4
2.5 5.0 7.5
τ (ms)
PDF (arb. u.)
0.25
0.00
τ=1.4
2.5 5.0 7.5
τ (ms)
PDF (arb. u.)
0.5
0.0
τ=1.4
2.5 5.0 7.5
τ (ms)
PDF (arb. u.)
0.25
0.00
τ=1.4
2.5 5.0 7.5
τ (ms)
PDF (arb. u.)
0.25
0.00
τ=1.4
2.5 5.0 7.5
τ (ms)
PDF (arb. u.)
0.5
0.0
τ=1.4
2.5 5.0 7.5
τ (ms)
PDF (arb. u.)
0.25
0.00
τ=1.3
2.5 5.0 7.5
τ (ms)
PDF (arb. u.)
0.25
0.00
τ=1.3
2.5 5.0 7.5
τ (ms)

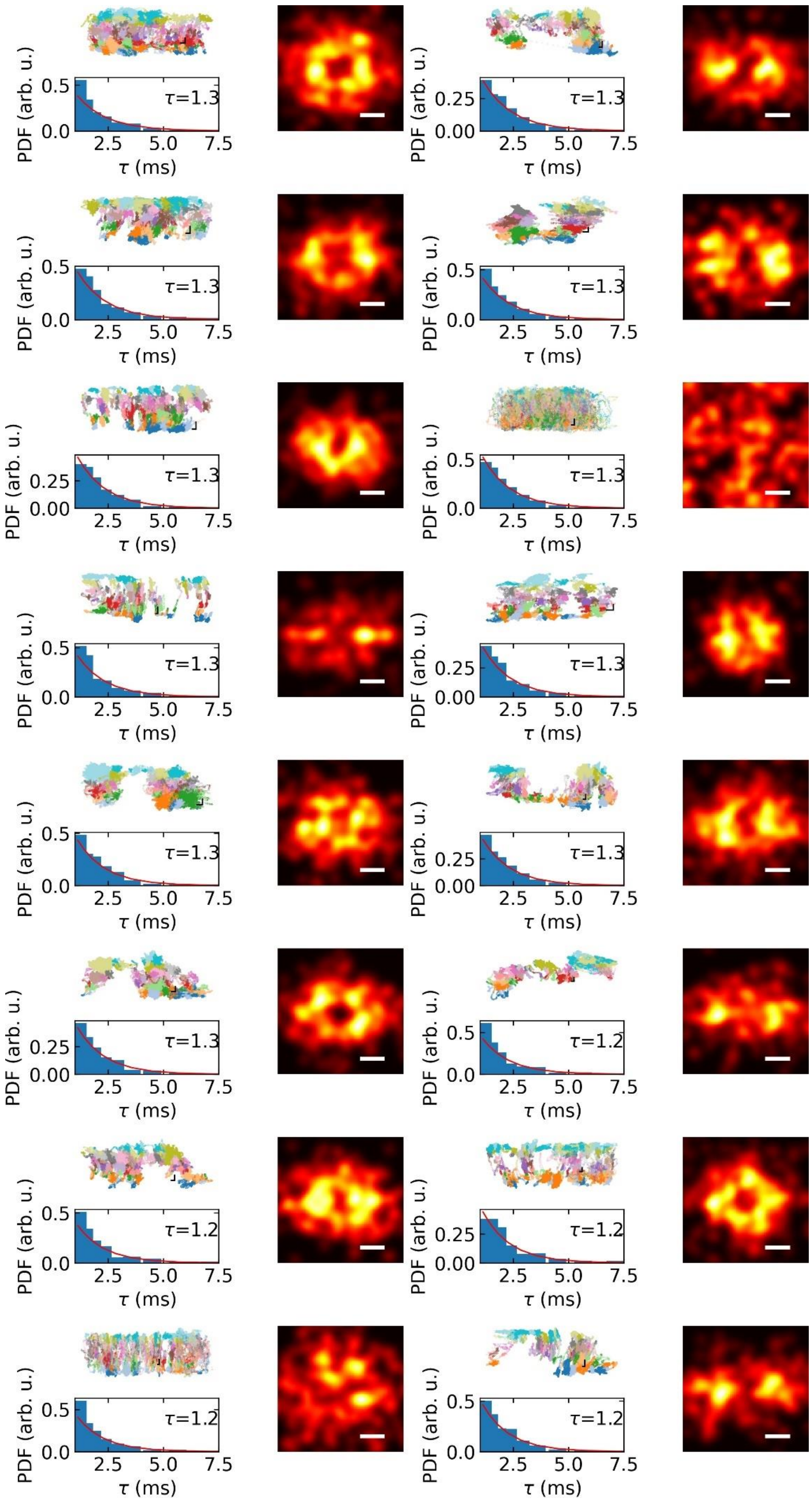
PDF (arb. u.)
0.5
0.0
τ=1.3
2.5 5.0 7.5
τ (ms)
PDF (arb. u.)
0.25
0.00
τ=1.3
2.5 5.0 7.5
τ (ms)
PDF (arb. u.)
0.5
0.0
τ=1.3
2.5 5.0 7.5
τ (ms)
PDF (arb. u.)
0.5
0.0
τ=1.3
2.5 5.0 7.5
τ (ms)
PDF (arb. u.)
0.25
0.00
τ=1.3
2.5 5.0 7.5
τ (ms)
PDF (arb. u.)
0.5
0.0
τ=1.3
2.5 5.0 7.5
τ (ms)
PDF (arb. u.)
0.5
0.0
τ=1.3
2.5 5.0 7.5
τ (ms)
PDF (arb. u.)
0.25
0.00
τ=1.3
2.5 5.0 7.5
τ (ms)
PDF (arb. u.)
0.5
0.0
τ=1.3
2.5 5.0 7.5
τ (ms)
PDF (arb. u.)
0.25
0.00
τ=1.3
2.5 5.0 7.5
τ (ms)
PDF (arb. u.)
0.25
0.00
τ=1.3
2.5 5.0 7.5
τ (ms)
PDF (arb. u.)
0.5
0.0
τ=1.2
2.5 5.0 7.5
τ (ms)
PDF (arb. u.)
0.5
0.0
τ=1.2
2.5 5.0 7.5
τ (ms)
PDF (arb. u.)
0.25
0.00
τ=1.2
2.5 5.0 7.5
τ (ms)
PDF (arb. u.)
0.5
0.0
τ=1.2
2.5 5.0 7.5
τ (ms)
PDF (arb. u.)
0.5
0.0
τ=1.2
2.5 5.0 7.5
τ (ms)

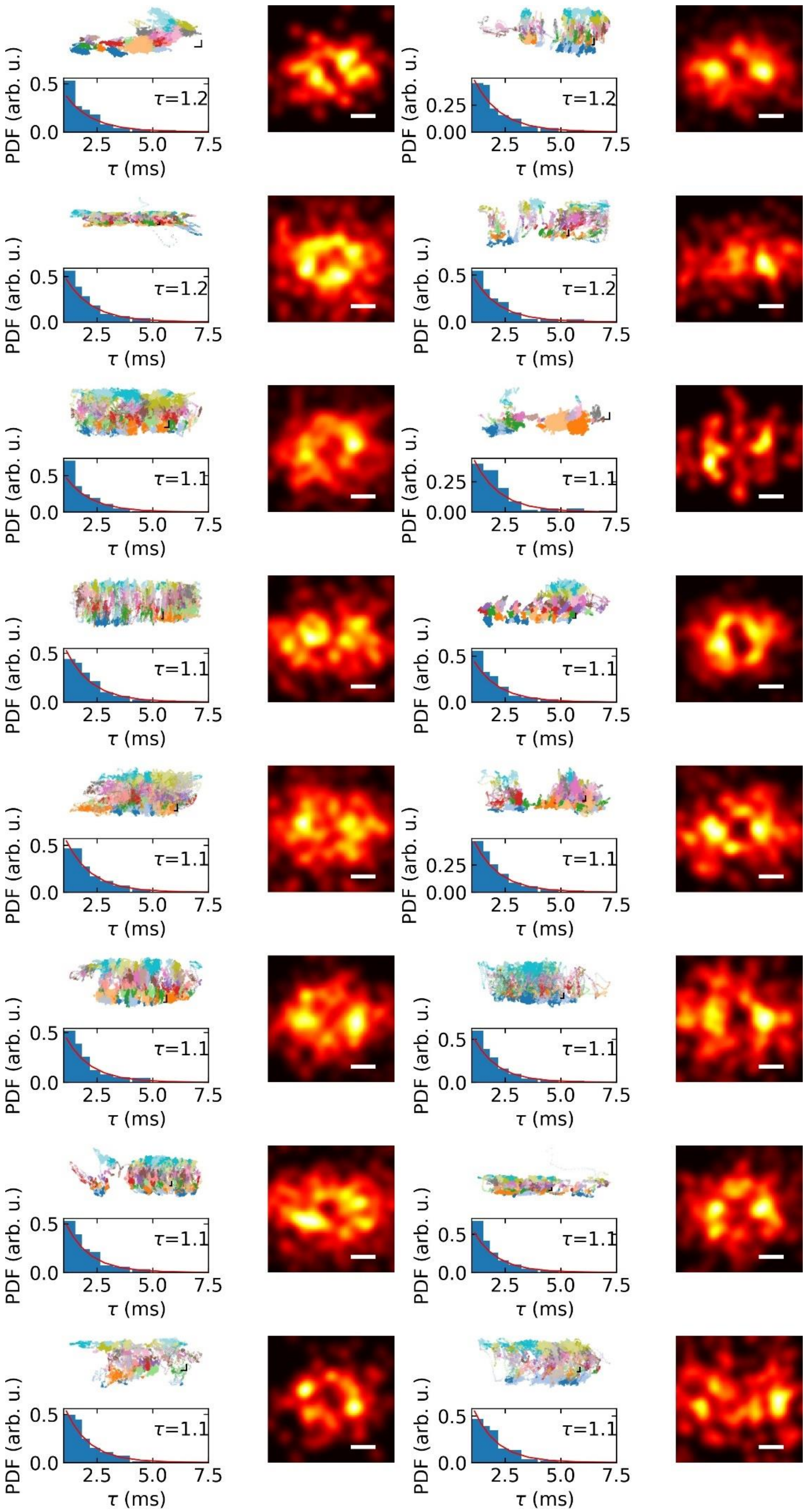
PDF (arb. u.)
0.5
0.0
τ=1.2
2.5
5.0
7.5
τ (ms)
PDF (arb. u.)
0.25
0.00
τ=1.2
2.5
5.0
7.5
τ (ms)
PDF (arb. u.)
0.5
0.0
τ=1.2
2.5
5.0
7.5
τ (ms)
PDF (arb. u.)
0.5
0.0
τ=1.2
2.5
5.0
7.5
τ (ms)
PDF (arb. u.)
0.5
0.0
τ=1.1
2.5
5.0
7.5
τ (ms)
PDF (arb. u.)
0.25
0.00
τ=1.1
2.5
5.0
7.5
τ (ms)
PDF (arb. u.)
0.5
0.0
τ=1.1
2.5
5.0
7.5
τ (ms)
PDF (arb. u.)
0.5
0.0
τ=1.1
2.5
5.0
7.5
τ (ms)
PDF (arb. u.)
0.5
0.0
τ=1.1
2.5
5.0
7.5
τ (ms)
PDF (arb. u.)
0.25
0.00
τ=1.1
2.5
5.0
7.5
τ (ms)
PDF (arb. u.)
0.5
0.0
τ=1.1
2.5
5.0
7.5
τ (ms)
PDF (arb. u.)
0.5
0.0
τ=1.1
2.5
5.0
7.5
τ (ms)
PDF (arb. u.)
0.5
0.0
τ=1.1
2.5
5.0
7.5
τ (ms)
PDF (arb. u.)
0.5
0.0
τ=1.1
2.5
5.0
7.5
τ (ms)
PDF (arb. u.)
0.5
0.0
τ=1.1
2.5
5.0
7.5
τ (ms)
PDF (arb. u.)
0.5
0.0
τ=1.1
2.5
5.0
7.5
τ (ms)

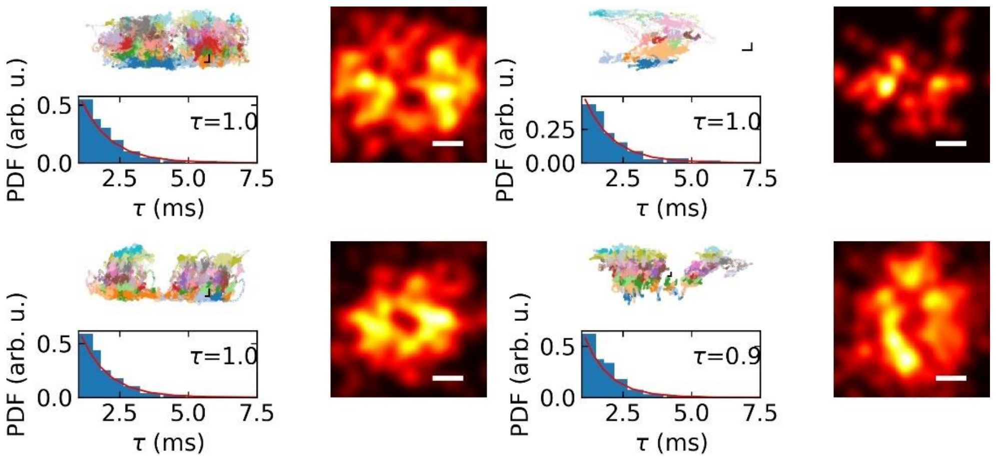


Figure S6. All the step detection figure for all the traces (Figure 5–type of analysis)

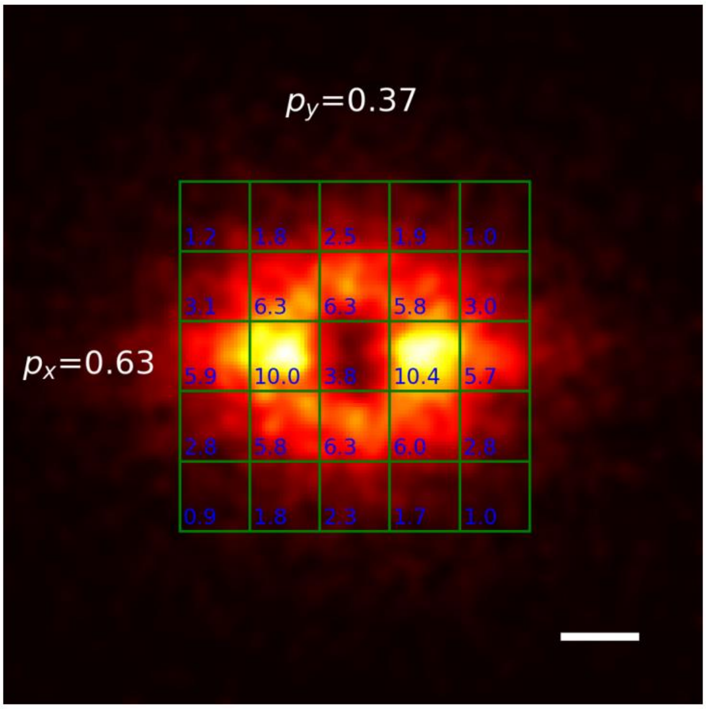


Figure S7. Accumulated 2D histogram of the displacement vector from all measured trajectories. The green grid indicates the grid of tubulin dimers in the MT lattice. Probabilities in percentages of given step size are written in each tubulin square.

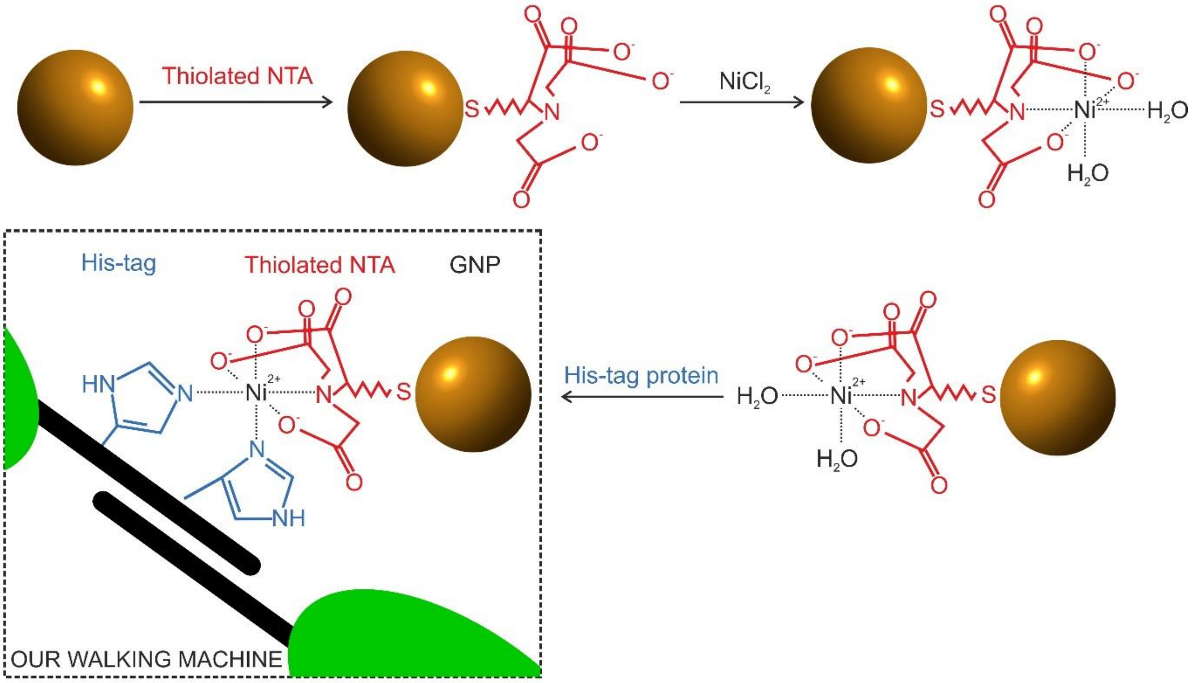


Figure S8. Illustration of the GNP attachment to the Ase1 molecule.

### Supplementary Video captions

**Supplementary Video S1. Visualization of the 3D trajectory of Ase1-GNP on MT.** Time evolution of one Ase1 trajectory corresponding to the data shown in Fig 2a. Static blue dots indicate a projection of the tubulin lattice. Scale bars are 20nm; previewed at 15x slow motion; *t* indicates the measurement frame time, the last 2ms (90 localizations) are color-coded separately.
